# Short-range interactions govern the dynamics and functions of microbial communities

**DOI:** 10.1101/530584

**Authors:** A. Dal Co, S. van Vliet, D. J. Kiviet, S. Schlegel, M. Ackermann

## Abstract

Communities of interacting microbes play important roles across all habitats on earth. These communities typically consist of a large number of species that perform different metabolic processes. The functions of microbial communities ultimately emerge from interactions between these different microbes. In order to understand the dynamics and functions of microbial communities, we thus need to know the nature and strength of these interactions. Here, we quantified the interaction strength between individual cells in microbial communities. We worked with synthetic communities of *Escherichia coli* bacteria that exchange metabolites in order to grow. We combined single-cell growth rate measurements with mathematical modeling to quantify metabolic interactions between individual cells and to map the spatial interaction network within these communities. We found that cells only interact with other cells in their immediate neighborhood. This short interaction range limits the coupling between different species and reduces their ability to perform metabolic processes collectively. Our experiments and models demonstrate that the spatial scale of biotic interaction plays a fundamental role in shaping the ecological dynamics of communities and the functioning of ecosystems.

## Inrro

Biological interactions are pervasive in nature, where organisms across all domains of life are connected through dense interaction networks^1^. These interactions influence what individual organisms do - the expression of their phenotypic traits and their rates of growth, reproduction and survival. The effects of interactions on individual organisms scale up to determine the dynamics and functions of ecosystems^2^. In natural systems, these interactions often emerge in spatially structured settings, where individuals interact preferentially with other individuals that are close in space^3^. To understand and predict the properties of such structured communities, we thus first need to understand the spatial interaction network between individual organisms, that is, understand the nature and strength of interactions between individuals as a function of their spatial position in a community. Then, we need to understand how these interactions scale up to give rise to processes at the community level^4^.

Our goal here was to analyze how local interactions in spatially structured communities determine community functions and dynamics. We focused on communities of interacting microbes. Microbial communities play important roles in all habitats on our planet. For example, microbial communities in the environment drive the global cycling of elements^5^, while the microbial community in our gut affects our physiology, cognition and emotion^6^. These community functions are based on biotic interactions between species. Microbial communities typically consist of hundreds to thousands of different microbial species that interact with each other in numerous ways^7^. These interactions are often based on diffusion-mediated exchange of molecules between cells^8–10^. Many microorganisms are unable to synthesize all the cellular building blocks required to grow and thus take up metabolites released by other cells^11–13;^ moreover, microorganisms often consume resources partially and exchange metabolic intermediates with other cells^14, 15^; finally, many microorganisms exchange signaling molecules with other cells to coordinate their activities^16^.

Most of these microbial interactions arise in a spatially structured situation. The majority of microorganisms across all habitats grows in biofilms, which are genetically diverse surface-associated communities embedded in a extracellular polymeric matrix^17^. In such spatially structured communities, one expects that the strength of the interaction between two organisms – that is, between two individual microbial cells – declines with increasing distance between them. A number of studies have predicted or observed that the strength of these interactions decays with the distance between cells^14, 18–22^. For example, mathematical models predict that yeast strains can exchange cellular building blocks across a range of about 100 µm, and experiments revealed that this influences the spatial self-organization of simple communities composed of such strains^18, 23^. In general, when the spatial range across which cells interact is small, the spatial arrangement of different cell types determines which cells interact with each other. Therefore, the interaction range between cells can strongly affect the collective functions and the dynamics of communities^8, 21, 24–26^. The interaction range is often an arbitrary parameter in theoretical models^24, 25^ or is experimentally measured in a heuristic and system specific way that cannot be easily generalized^19, 27^. We lack direct measurements of the interaction range between individual cells and a mechanistic understanding of the factors that determine this spatial scale. Progress in this direction would allow to build a general framework to predict which ecological interactions emerge in microbial communities and to understand how these interactions shape community properties.

Our aim here was to develop such a general framework. More specifically, our first main goal was to directly measure the interaction range in assembled microbial communities. Our second main goal was to obtain a mechanistic understanding of the factors that determine the interaction range in order to predict the interaction range in other systems. Our third goal was to assess the consequences of this interaction range at the level of the community. We combined time-resolved quantitative single cell measurements in a spatially structured synthetic community with mathematical modeling to address these goals.

## Results

We focused on a scenario where two bacterial genotypes exchange cellular building blocks that are essential for growth, a situation that is widespread in natural microbial communities^11, 28^. In such a situation, one expects that the interaction range between cells will strongly affect the growth of individuals and communities. This can be illustrated with a simple simulation of the cellular dynamics in a system composed of two interacting partner species (Fig. 1a). The simulation rests on three general assumptions: first, each cell can only receive compounds from cells belonging to the partner species that reside within the interaction range; second, the growth of individual cells depends on the fraction of the cells of the partner species within the interaction range; third, if a cell divides, it places an offspring on a neighbouring site. This simple simulation reveals that the average growth rate of individual cells is low when the interaction range is small (Fig. 1b). This key finding originates from a simple mechanism: most individuals are surrounded by their offspring; if they interact on a small spatial scale, they mostly interact with these offspring from which they cannot obtain the cellular building blocks they need. This finding is consistent with previous theoretical studies^24, 25^. This effect becomes stronger when organisms depend on compounds from two or more other species: a small interaction range reduces the probability that an individual is close enough to all of these partners simultaneously (Fig. 1b). The simulation shows that short-range interactions can reduce the growth rate of cells whenever these cells need to exchange compounds with other genotypes in order to grow, in line with previous observations^29^.

**Fig. 1:**
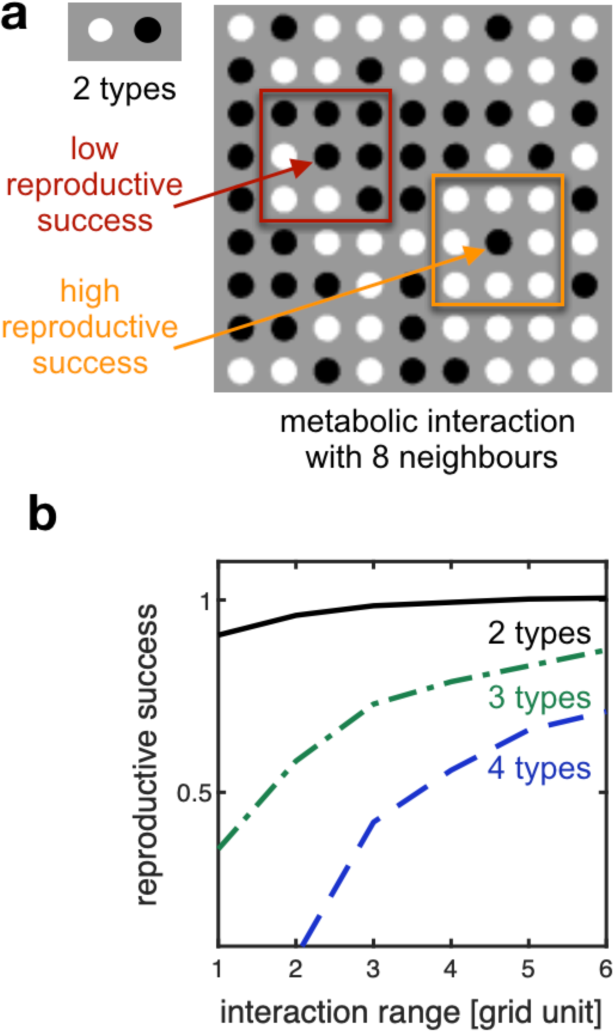
Interacting locally lowers reproductive success. **a** A grid is populated by two types of cells (black or white dots). The two types exchange compounds to reproduce, and place offspring on adjacent sites. The reproductive success of an individual increases linearly with the fraction of the partner cells within their interaction range. In the example shown here, the interaction range is one grid unit. **b** The reproductive success of individuals is lower in consortia with smaller interaction range. This decrease is already visible in consortia composed of two interacting types (as in panel a). It becomes more pronounced in consortia composed of three or four types, where an individual cell can only grow if all the other types reside in the individual’s interaction range.

### Cells in dense microbial communities interact on a range of a few cell lengths

Our first major goal was to quantify the interaction range experimentally. We constructed a microfluidic device for growing cells in monolayer communities and developed an analytical method to extract time-resolved quantitative single-cell data (Fig. 2). We focused on a synthetic consortium composed of two auxotrophic *Escherichia coli* strains. The first strain is unable to produce the amino acid proline and the second strain is unable to produce the amino acid tryptophan (Fig. 2a). Because cells naturally leak out amino acids, the two auxotrophs can grow together by exchanging the two amino acids through diffusion^28, 30, 31^. We grew our consortia in the microfluidic device and used automated image analysis to identify and track single cells so that we could measure their growth rate (Fig. 2b, Sup. Video S1). The goal of this analysis was to determine the spatial range from which a single cell could retrieve amino acids, that is, their interaction range. How fast an individual cell grows is expected to depend on the amount of the amino acid received, and this in turn depends on the number of partner cells inside the interaction range. In order to determine the size of the interaction range, we thus looked for the spatial range whose cellular composition best predicted the growth rate of individual cells (Fig. 2c, Sup. Video S2). Therefore, we measured the fraction *f_d_* of the partner within a distance *d* from a cell and determined the correlation between this fraction and the cells’ growth rate, for a large number of individual cells. The interaction range is then defined as the distance *d* where this correlation is maximal (Fig. 2d).

**Fig. 2:**
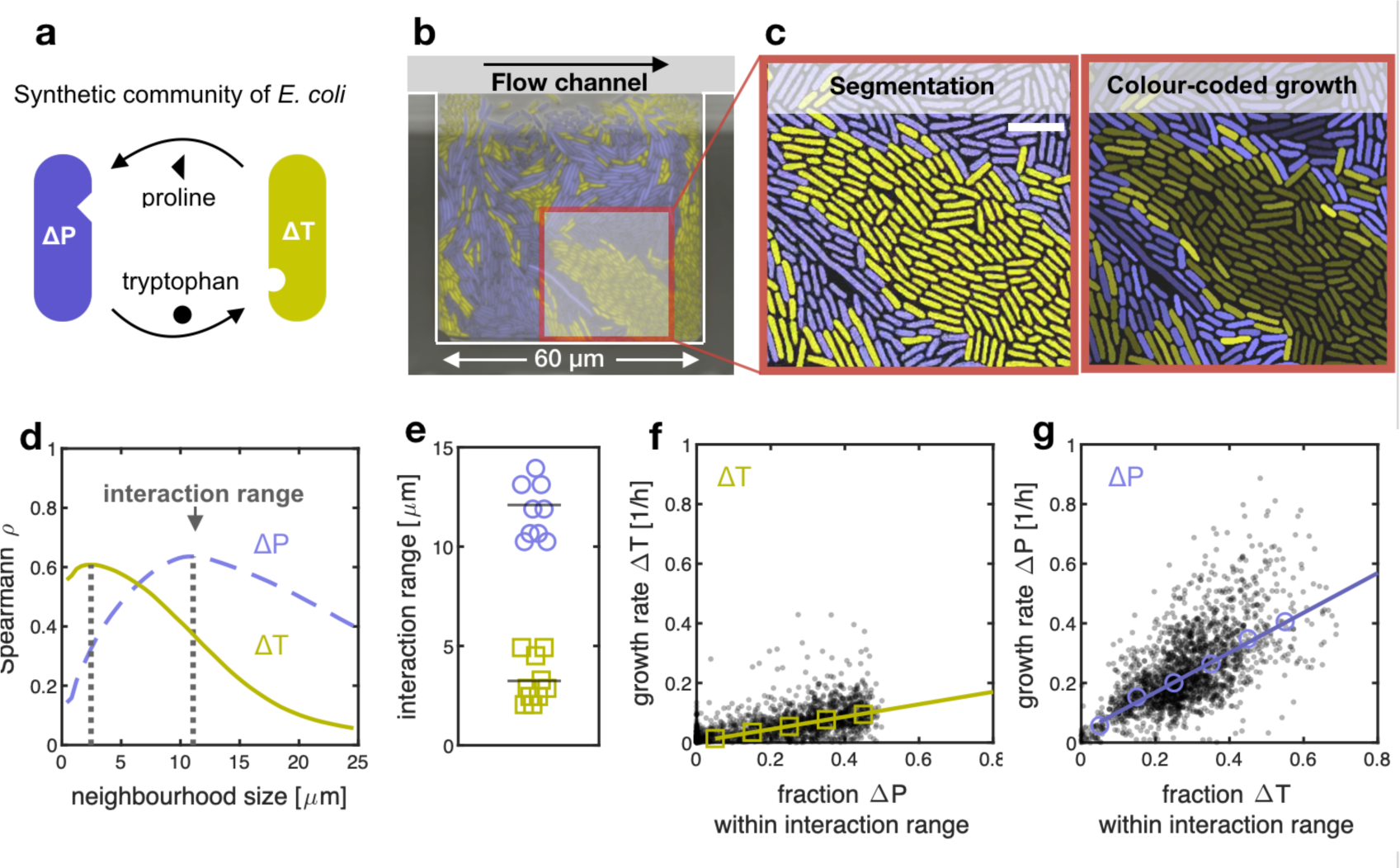
Auxotrophic cells that exchange cellular building blocks interact on a small range. **a** Synthetic communities of two auxotrophic strain of *E. coli* that depend on each other and that are labeled with constitutively expressed fluorescent proteins (here depicted as yellow and purple). **b** False colour image of a microfluidic chamber where cells grow in a monolayer. Continuous flow of culture media at the top of the chamber removes cells as soon as they are pushed out of the chamber. **c** Left: cells are segmented and tracked. Right: Cells are colour-coded based on their individual growth rate, with brighter colours indicating higher growth rates. Cells that are surrounded by the partner grow faster than cells that are surrounded by their own type. White scale bar 5 µm. **d** We calculated the correlation coefficients between the growth rates of individual cells and the fraction of the partner in a given neighbourhood size. When we plot the correlation coefficient as a function of the neighbourhood size, we observe that the strength of the correlation is maximal for an intermediate neighbourhood size (marked by dashed lines); we call this neighbourhood size the interaction range. **e** The two auxotrophs have different interaction ranges (10 biological replicates, ∼10,500 cells total). **f-g** Both auxotrophs grow faster with increasing fraction of the partner within the interaction range. Tryptophan auxotrophs (ΔT, f) generally have lower growth rates than proline auxotrophs (ΔP, g), as shown by the slopes of the linear regression (0.75 for ΔP and 0.21 for ΔT). Black dots: single cells (1,985 ΔP and 1,769 ΔT cells); Open symbols: binned median values; lines: linear regression on binned values.

This analysis revealed that the interaction range is on the order of only a few cell lengths (Fig. 2d). This is found consistently across ten biological replicates (∼10,500 cells analysed in total). Specifically, the interaction range of the tryptophan auxotroph cells is 3.2±0.4 µm (mean ± standard error of the mean), while the interaction range of the proline auxotroph cells is significantly larger at 12.1±0.5 µm (*p*<10^-5^, paired t-test, *n*=10, Fig. 2e). In other words, these cells live in a small world: they only interact with a small group of individuals around them. Cells can only grow well if their partner is among these individuals (Fig. 2f-g). In control experiments where amino acids were provided with the growth medium, the growth rate of individual cells does not depend on the proximity to the partner (Fig. S1).

### A mathematical model offers a mechanistic explanation for the small interaction range

Our second major goal was to obtain a mechanistic understanding of the factors that determine the interaction range. Why do cells only interact across such a small spatial range? We addressed this question with an individual-based model (Fig. 3a), where cells occupy a site on a grid with size 40×40. At every grid site, we describe the internal and external concentration of the two exchanged amino acids with a set of differential equations (see Fig. 3b and Methods for details). We assumed that the growth rate of auxotrophic cells is limited by the amino acid that they need, while they produce enough of the other amino acid so that it does not limit their own growth. Cells take up amino acids actively and leak them passively in the environment, where they diffuse. All model parameters were taken from literature, or were directly measured, apart from the two leakage rates, which were estimated from the data (Supplementary Information 3.4.2).

**Fig. 3:**
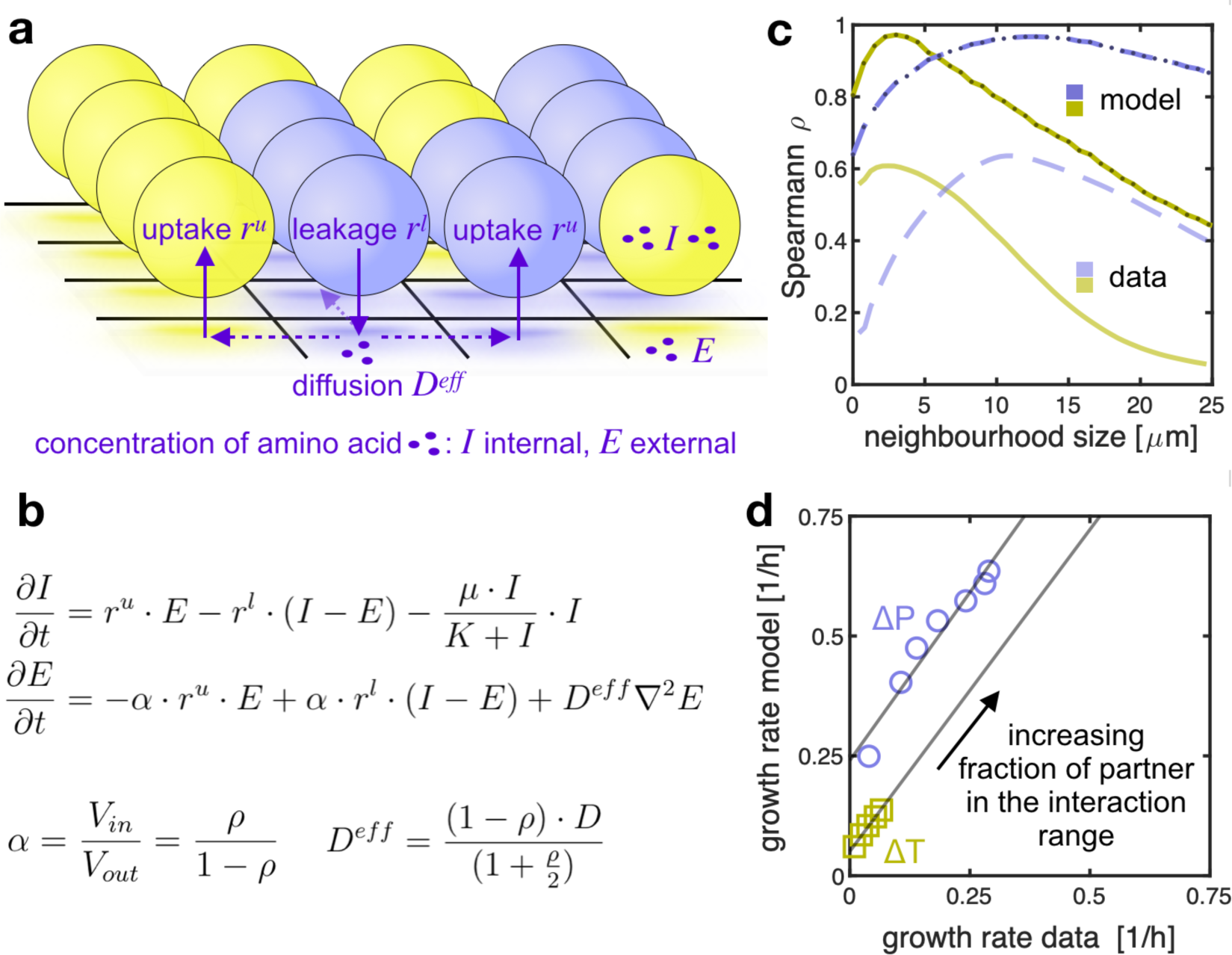
Mathematical model reveals mechanism of local interactions. **a** Individual-based model where amino acids are passively leaked, diffuse in the environment and are actively taken up. **b** At every grid site, we describe the internal (*I*) and external (*E)* concentration of amino acids with a set of differential equations (here shown for one amino acid only). These equations assume that the growth rate of auxotrophic cells is limited by the amino acid that they need, that cells take up amino acids actively (with rate *r^u^*) and leak them passively into the environment (with rate *r^l^*), where they diffuse. The effective diffusion constant (*D^eff^*) is lower when the density of cells *ρ* is higher. **c** The correlation analysis based on the model (dark curves) matches the results obtained from experimental data (light curves, identical to Fig. 2d). The predicted interaction range for ΔP is 12.8 µm (compared to 12.5 µm in the experimental measurements, *p*=0.25, t-test) and for ΔT is 3.0 µm (compared to 3.2 µm in the experimental measurements, *p*=0.51, t-test). **d** The predicted and experimentally measured growth rates are strongly correlated (*r^2^*=0.95 for ΔP and *r^2^*=0.99 for ΔT, Pearson correlation). We grouped cells based on the fraction of the complementary partner in their interaction range and for each group we compared the measured growth rate (x-axis, same data as Fig. 2f-2g) to the predicted growth rates (y-axis). Each symbol represents a single group.

We first tested if our model can predict the interaction range that we experimentally measured. For that, we applied the model to our measured spatial arrangements of the two cell types, calculated the concentration of amino acids in space by solving the equations at steady state, and subsequently calculated the theoretical growth rates of individual cells from the local concentration of these amino acids. Then we estimated the interaction range by correlating the theoretical growth rates of cells with the fraction of their partner in their neighbourhood, as we did with the experimentally measured growth rates. The interaction range we found deviated less than 7% from the experimental interaction range (Fig. 3c). Our model thus predicts the interaction ranges that we measured experimentally. Moreover, our model predicts that the growth rate of the auxotrophs increases with the fraction of the partner within the interaction range, in agreement with the experimental data (Fig. 3d). We conclude that our model is consistent with the experimental data: the mechanisms of amino acid exchange we propose can explain how cells interact in these communities.

### The interaction range is set by few key parameters

Our model reveals that the short interaction range that we measured is mainly a consequence of high uptake rates of amino acids and dense packing of cells. This becomes evident when we look at a second length scale, which is directly proportional to the interaction range: the *growth range*, the length scale describing the decrease in growth away from a straight interface separating the two cell types (see Fig. 4a). When the two cell types are in such a symmetric configuration, the concentration of amino acids can be approximated analytically, and from this the growth of the two cell types can be predicted (Supplementary Information 3.4.3). The growth range of each type in this symmetric configuration is proportional to its interaction range in any complex spatial configuration (Fig. 4b). The analytical expression of the growth range of each auxotroph is (see Fig. S9 for comparison with the numerical solution):

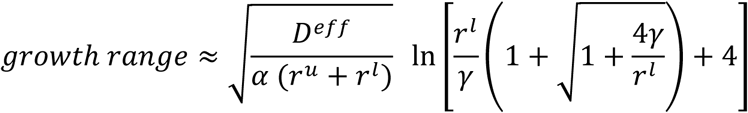

where *r^l^* and *r^u^* are the leakage and the uptake rates of the amino acid each auxotrophs needs, 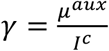 where *I*^*c*^ is the constant internal concentration of the amino acid in producing cells and *μ*^*aux*^ the maximal growth rate of each auxotroph (equal to the growth rate of the wild type in our case; see Fig. S7). *D^eff^* is the effective diffusion constant which accounts for the density of cells, and *α* the ratio between the volume of intra- and extracellular environment; the ratio of these parameters depends on the density *ρ* of cells:

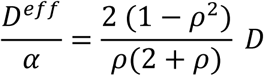

**Fig. 4:**
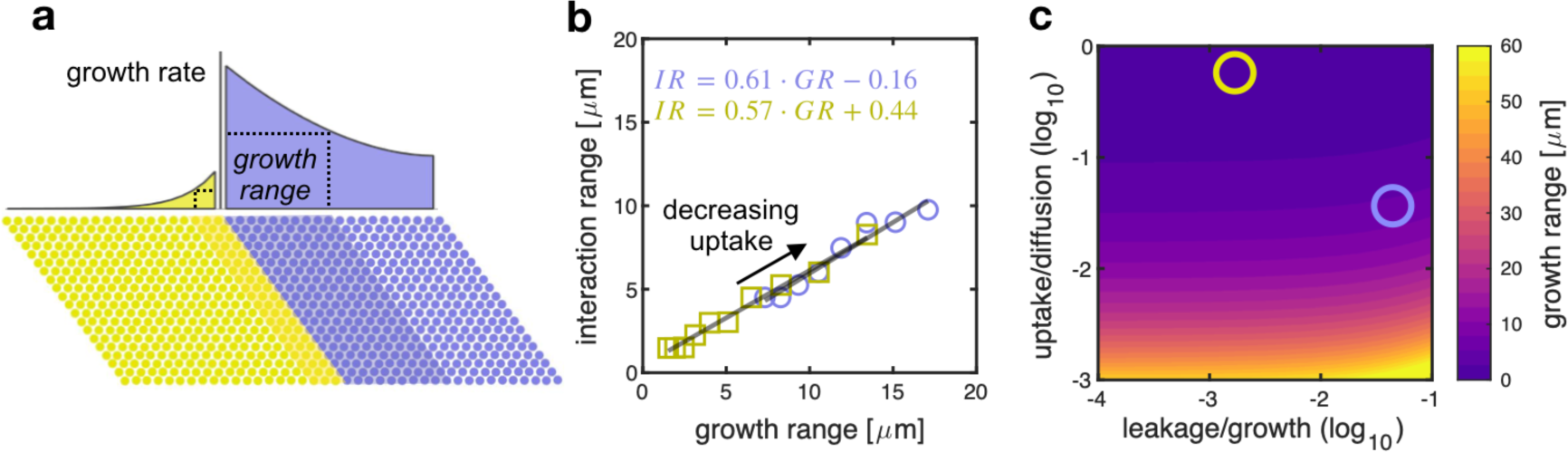
Interaction range is small when uptake rates are high. **a** For a symmetric arrangement with a straight interface between the two types, we can analytically calculate the region (shaded) in which the cellular growth rate is at least half of the maximal growth rate observed at the interface; we call this region the growth range. The growth range can be calculated from biochemical parameters. **b** The growth range is proportional to the interaction range (*r^2^*=0.95 for ΔP and *r^2^*=0.99 for ΔT, Pearson correlation). When we decrease, in the model, the rate at which cells take up amino acids, the growth range and the interaction range increase; each open symbols shows the growth range and interaction ranges calculated using our model for different values of the uptake rate of amino acids. The growth range is calculated analytically while the interaction range is estimated numerically. **c** Growth range as a function of leakage and uptake rate (relative to maximum growth rate and diffusion constant respectively). Growth range (and interaction range) of the tryptophan auxotroph (yellow circle) and proline auxotroph (purple circle) are small because the ratio between uptake and diffusion is high.

From the mathematical expression, we see that the growth range (and thus the interaction range) depends on the uptake, leakage and diffusion rates of the amino acids and on the density of cells. Specifically, the growth range (and the interaction range) is small in consortia where the leakage rate is low, the uptake rate of the exchanged compounds is high compared to their diffusion (Fig. 4c), and the density of cells is high. High cell densities reduce the interaction range by reducing effective diffusion (Fig. S10). This means that denser cellular aggregates tend to have cell-cell interactions that are more localized, and therefore cellular density is an important parameter modulating interactions in these aggregates^32–34^. While cell density alters the interaction range of different types in a consortium in the same way, the other parameters modulate the interaction range of each cell type separately. The difference in interaction range between types depends on the difference in uptake, leakage and diffusion constants of the amino acids they exchange. Diffusion constants typically vary only over a small range between different amino acids and can thus not explain large differences in the interaction ranges. However, uptake and leakage rates can vary substantially between amino acids (for example the ratio between the diffusion constants of tryptophan and proline 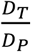 is 0.75, while the ratio between the rates at which the two amino acids are taken up, 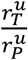, is 12). From the analytical expression of the growth range, we can show that the growth range (and the interaction range) depends more strongly on the uptake rate than on the leakage rate (Fig. 4c and Fig. S8). While leakage rates have a minor effect of the growth range (*how far* a cell can grow away from the partner), they have a major effect on the growth rates of cells (*how fast* cells grow). More precisely, we can show that leakage rates set the maximum growth rate *μ^max^* that an auxotroph can reach when fully surrounded by the partner (Supplementary Information 3.4.1):

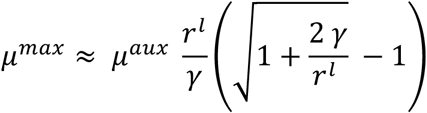

The growth range and *μ^max^* vary independently, being primarily modulated by the uptake rates and the leakage rates, respectively. Our findings apply generally to any dense microbial assembly where molecules are exchanged by leakage, diffusion and uptake. For example, the same model could estimate the length scale of cell-cell communication via molecules that are taken up or degraded by the recipient, such as quorum-sensing molecules.

### Interaction range and patch size are two different length scales

According to our mechanistic model, the interaction range represents the size of the neighbourhood from which cells can retrieve the amino acids produced by the partner. It might seem intuitive to measure this neighbourhood using a simpler quantifications such as the patch size of the different genotypes^35^. We tested the validity of this method and found that interaction range and patch size are potentially two different length scales: we verified that the interaction range of two auxotrophs varies less than 50% for the range of patch sizes observed in our chambers (a small change compared to the fourfold difference in interaction range between the auxotrophs, Fig S3). The reason is that the interaction range is set by the cell density and few biochemical parameters (uptake relative to diffusion), while patch size depends mostly on the physics of cell division and movement. In general, we see no significant correlation between size of patches and average growth in these patches (Fig. S4).

### A small interaction range affects growth and dynamics of the whole community

How do short-range interactions between individual cells affect community-level dynamics? Our communities show consistent dynamics in time: within about 25 hours, all 61 replicate communities reach a steady state composition, and in all but two of the communities the tryptophan auxotroph is in minority (median fraction of total biomass = 0.23, Fig. 5a). This shift in the composition of the community arises from the individual-level properties that we measured: the proline auxotroph tends to increase in frequency because of a double advantage: it has a higher *μ^max^* and a larger interaction range than the tryptophan auxotroph; as a consequence, the growth rate of the proline auxotroph is less sensitive to the spatial arrangement. The differences in interaction range and *μ^max^* drive the community to its equilibrium composition where the tryptophan auxotroph is in minority.

**Fig. 5:**
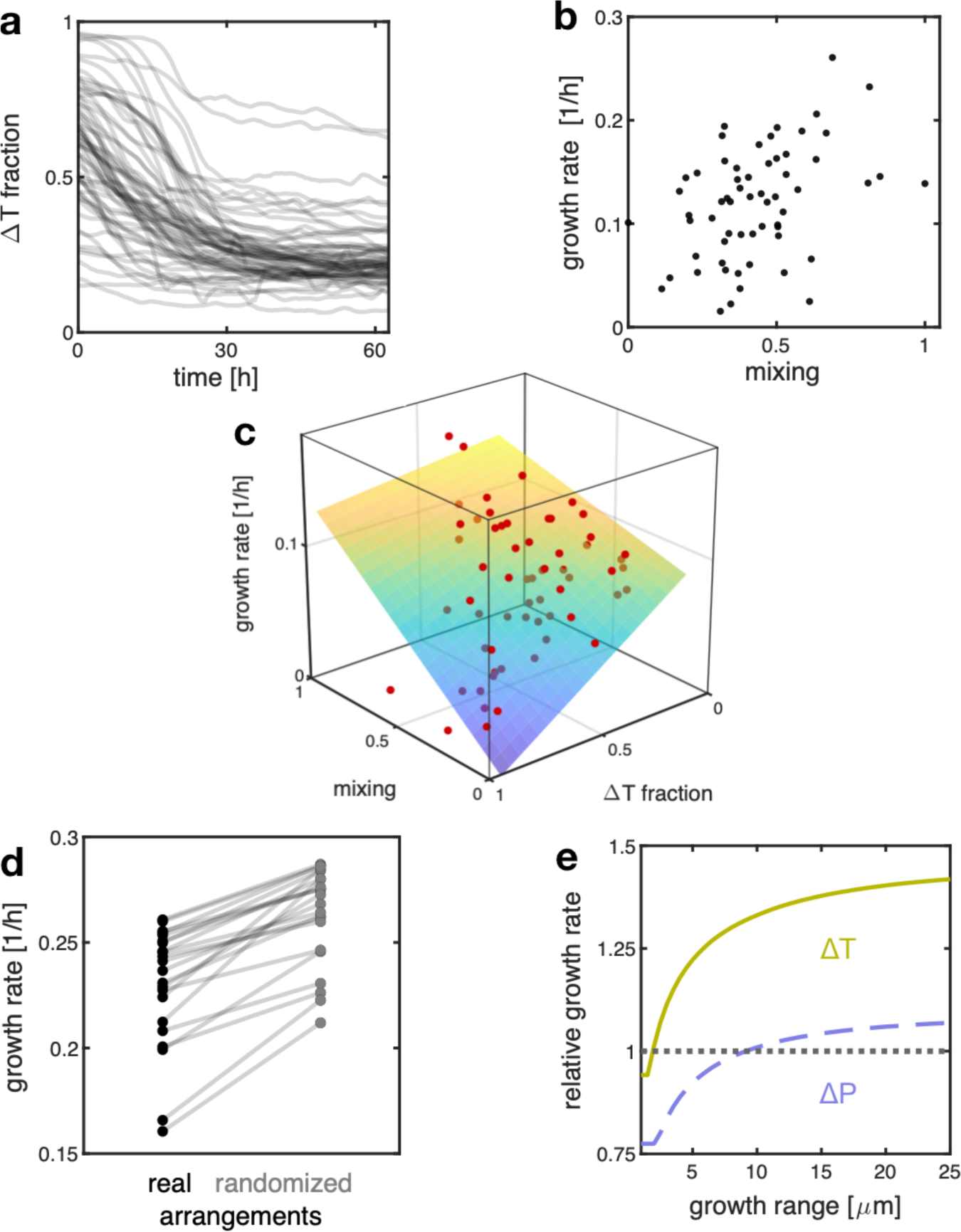
The small interaction range between cells reduces productivity of the consortium. **a** Communities equilibrate at compositions of 23% (median, n=61) of the tryptophan auxotrophs; deviations in two replicates are due to large clusters of tryptophan auxotrophs in the back of the chamber. **b**-**c** Growth of the communities increases with the mixing of the two cell types (partial correlation analysis with fraction as control variable Spearman ρ = 0.44, p<10^-3^, n=61). Each dot shows growth and mixing of one community. In panel **c**, also the fraction of tryptophan auxotroph is shown; the plane depicts the multiple linear regression of the growth rate on the mixing and the fraction of tryptophan auxotroph (the colour of the surface indicates the growth rate); data points above the plane are shown in bright red, while data points below the plane are shown in dim red)**. d** Randomizing community arrangements leads to higher mixing and higher predicted average growth rates of individuals (number of randomizations = 20, p<10^-5^, paired t-test, n=22). This indicates that the unmixing of the two types (which is removed by randomizing the arrangements) decreases average growth rates. **e** The model predicts an increase in the average growth rate (relative growth above one), when the growth range (and thus interaction range, see Fig. 4b) increases. Here we simulated a closed system, where amino acids are not lost from the system through diffusion; the growth range was varied by changing the uptake rate of amino acids in the model.

A final and important question is whether the small interaction range between cells limits the growth of the community as a whole. This question brings us back to our central hypothesis, that a small interaction range limits the exchange of resources and hinders collective metabolism because many cells reside within groups of their own type. We therefore tested if communities with higher level of mixing of the two cell types grew faster. The average growth rate of cells in a community is determined by several factors, including the proportion of the two types and their level of mixing. We measured growth of the 61 communities after 16 hours using an image analysis method based on optical flow, which provides an estimate of the average growth rate of cells in each chamber. We found that communities grew faster when they had a higher level of mixing of the two types (Fig. 5b-c).

We further tested the effect of mixing on the average growth of our communities using our model. Specifically, we tested the prediction of our simple cellular automaton, that cells in our communities would grow faster if the interaction range was larger or if the spatial arrangement was more mixed. We tested these predictions by applying our model to experimentally observed and computationally altered spatial arrangements. Specifically, we randomized the observed spatial arrangements to disrupt kin clusters and we found that the average predicted growth rate of individuals increases (Fig. 5d). Likewise, if we simulate a closed system (corresponding for example to a large biofilm) where no amino acids are lost from the community, we find that lowering the uptake rates of amino acids and thereby increasing the interaction range leads to an increase of the average predicted growth rate of individuals (Fig. 5e). However, there is a tradeoff: in systems that are open and where metabolites can diffuse away from the cells (like our chambers), very low uptake rates can also reduce the average growth rate because of diffusional loss (Fig. S5).

## Discussion

Here we developed a method to directly measure the interaction range that can be applied to a large number of microbial systems. We focused on a synthetic community of two genotypes exchanging amino acids and we found that the cells in our community interact on a short range and that this lowers their growth rates. In general, we expect the interaction range to fundamentally affect the functioning of any assembly of interacting microorganisms. The specific effects will depend on the nature of the interactions. For example, short-range interactions can stabilize the cooperative production of molecules, as they ensure that these molecules are only accessible to cells that also contribute to production, and are inaccessible to non-producing individuals^21, 36^. In contrast, short-range interactions generally can impede mutualistic cross-feeding^18^, although they can have a stabilizing effect by preventing ecological invasion by non-contributing mutants^23^.

The ecological and evolutionary outcome of cooperation and competition can change dramatically when interactions are limited to a small neighbourhood^8, 29, 37^ and therefore the interaction range is a crucial feature of any spatially structured ecological system. Here we found that the interaction range between individuals is on the order of a few cell lengths in a microbial assembly where the production of cellular building blocks is distributed across different cell types. We predict that the interaction range is generally small whenever the density of cells is high and the uptake of the molecules mediating the interaction is fast compared to their diffusion. We thus expect the interaction range to be small in dense assemblies where cells exchange cellular building blocks, signaling molecules or metabolites that bind^38^ or digest extracellular nutrients^39, 40^. Finally, we showed that, if interaction ranges are small, the spatial unmixing of cell types through local growth can hinder metabolic exchange between different cell types and reduce community growth.

Our work suggests that knowing at which spatial scale organisms interact is crucial for understanding the ecological dynamics and functions of communities. Here, we worked with microbial systems, where interspecies interactions are often based on the uptake and release of diffusible metabolites. In plant communities, the ecological dynamics are mostly shaped by competition for light and nutrients^41^ as well as by facilitation^42^. In communities of predators and prey, interactions are based on encounter rates and thus by the movements of individuals^43^. In all these cases, the interaction strength is expected to decline with distance between individuals. Understanding how local interactions scale up to determine the dynamics and functions of such spatially structured communities is thus a central goal.

## Methods

### Strains

All experiments were performed using strains derived from *E. coli* MG1655; these strains are ΔtrpC-GFP (MG1655 *trpC*::frt, PR-*sfGFP*), ΔtrpC-RFP (MG1655 *trpC*::frt, PR-*mCherry*), ΔproC-GFP (MG1655 *proC*::frt, PR-*sfGFP*), and ΔproC-RFP (MG1655 *proC*::frt, PR-*mCherry*). The ΔproC strains are unable to produce proline due to a deletion in *proC*, the ΔtrpC are unable to produce tryptophan due to a deletion in *trpC*^31^. The auxotrophic strains were made by transferring the respective kanamycin cassettes from the keio-collection^44^ into TB204 and TB205^45^ using lambda Red mediated recombination^46^. TB204 and TB205 are *E. coli* MG1655 derivatives that constitutively express sfGFP or mCherry from the lambda promoter (P_R_) from the chromosome. In brief, the kanamycin cassette including the homologous flanking regions were amplified by PCR from JW0377 (*proC*::*kan*) and JW1254 (*trpC*::*kan*)^44^ and transformed into TB204 and TB205 harbouring the pSim8 plasmid (kindly provided by Donald L. Court). Primer sequences used:

U_proC_fw: CAT AAA GTC ATC CTT TGT TGG G
D_proC_rv: CTT TAC GGA TTA GTG TGG GG
U_trpC_fw: AAC GTC GCC ATG TTA ATG CG
D_trpC_rv: GAA CTG AGC CTG AAA TTC AGG

The kanamycin cassette was transferred into a fresh strain of TB204 or TB205 using P1 mediated generalized transduction. Upon successful transduction, the phenotypes of the strains were confirmed (no growth without proline or tryptophan) and the kanamycin cassettes removed from the genome using the FLP-recombinase from plasmid pCP20^46^. We confirmed the ability of our two auxotrophs to grow together by receiving the amino acid they cannot produce from their partner, as reported in previous work^31^.

### Media and growth condition

Monocultures of the two auxotrophs strains were started from a single colony taken from a LB-agar plate and were grown overnight at 37°C in a shaker incubator in M9 medium (47.76 mM Na_2_HPO_4_, 22.04 mM KH_2_PO_4_, 8.56 mM NaCl and 18.69 mM NH_4_Cl) supplemented with 1mM MgSO_4_, 0.1 mM CaCl_2_, 0.2% glucose (all from Sigma-Aldrich), 50 µg/L of L-proline (434 mM) and 20 µg/L L-tryptophan (98 mM) and 0.1% Tween-20 (added to facilitate loading of cells in microfluidic device). Cells were loaded in stationary phase in a microfluidic device and grown in the same media. After approximately 10 hours, cells exited lag phase and started to fill the chambers. The medium was then switched to M9 medium + 0.2% glucose + 0.1% Tween-20 with no amino acids. This medium was fed for the whole duration of experiment (approximately three days). Imaging was started three hours before switching to this medium, to have a control of cellular growth with amino acids in the medium.

### Microfluidic experiment

The microfluidic devices consisted of chambers of 60×60 µm and 0.76 µm in height facing a feeding channel of 22 µm in height and 100 µm in width. Masks for photolithography were ordered at Compugraphics (Jena, Germany). The master mold was made on a silicon wafer, by applying SU8 photoresist in two steps (the first step to make the layer for the growth chambers and the second step to make the layer for the feeding channel). To make the chips used for the experiments, Polydimethylsiloxane (PDMS, Sylgard 184 Silicone Elastomer Kit, Dow Corning) was mixed in a ratio of 1.5:10 and poured on the dust-free master mold, degassed in a desiccator for 30 minutes, and baked for around one hour at 80°C for curing. PDMS chips of approximately 2 cm × 3.5 cm were cut out around the structures on the wafer. Holes for medium supply and outlet were punched (diameter of holes 1.2 mm). PDMS chips were bound to round (50 mm diameter) glass coverslips (Menzel-Gläser, Braunschweig, Germany) by treating them for 30 seconds at maximum power in a Plasma Cleaner (PDC-32G-2, Harrik Plasma, New York, USA), and left on a hated plate at 100°C for one minute for binding. Before an experiment, a small amount of medium was flushed into the channels using a pipette to wet the chambers. Then air was pushed through the main channel (medium remained in the chambers). Cells in stationary phase, from overnight culture (approximately 14 hours) were concentrated approximately 100 times by centrifugation (5,000×*g*, 5 min.) and loaded into the chip using a pipette. Cells were pushed in the side chambers with the help of small air bubbles flowing through the main channel. Once a sufficient number of cells were pushed inside the chambers, fresh medium was pumped through the flow channel. For all experiments, syringe pumps (NE-300, NewEra Pump Systems) with 50 ml syringes containing the medium were used. Tubing (Microbore Tygon S54HL, ID 0.76 mm, OD 2.29 mm, Fisher Scientific) was connected to the syringes using 20G needles (0.9 mm × 70 mm), which were directly inserted into the tubing. Smaller tubing (Teflon, ID 0.3 mm, OD 0.76 mm, Fisher Scientific) was then inserted into the bigger tubing and directly connected to the inlet holes in the PDMS chip. Medium switches were performed by disconnecting the bigger tubing from the syringe and reconnecting it to new syringes. All experiments were run at a flow rate of 0.5 ml/h. The flow rate is high enough that amino acids do not accumulate in the feeding channel and are not exchanged via the main channel. In fact no growth was observed in chambers hosting only one of the two auxotrophs during the whole duration of the experiment.

### Microscopy

Time-lapse microscopy was done using fully automated Olympus IX81 inverted microscopes (Olympus, Tokyo, Japan). Images were taken using a 100X NA1.3 oil objective (Olympus) with 1.6X manual auxiliary magnification and an ORCA-flash 4.0 v2 sCMOS camera (Hamamatsu, Hamamatsu, Japan). Fluorescent imaging was done using a X-Cite120 120 Watt high pressure metal halide arc lamp (Lumen Dynamics, Mississauga, Canada) and Chroma 49000 series fluorescent filter sets (N49002 for GFP and N49008 for RFP, Chroma, Bellows Falls, Vermont). Focus was maintained using the Olympus Z-drift compensation system and the entire setup was controlled with Olympus CellSens software. The sample was maintained at 37°C with a microscope incubator (Life imaging services, Basel, Switzerland). Several positions were imaged on the same microfluidic device and images were taken every ten minutes.

### Image analysis

All image processing was done using Matlab (version 2016A and newer, MathWorks, Natick, Massachusetts) and Vanellus software (written by D. J. Kiviet, accessible at https://github.com/daankiviet/vanellus). Time-lapse frames were first registered and cells were then segmented using customized segmentation algorithms. Two different algorithms for segmentation were used: the ‘*segmentation of biomass algorithm’* and the ‘*segmentation of cells algorithm’*. The ‘*segmentation of biomass algorithm’* identifies the green and red biomass in the chamber: images were first cropped along the profile of the microfluidic chambers (up to 8 µm from the outlet), and biomass was segmented on the phase contrast image and assigned to its relative colour after deconvolution; the algorithm was optimized to give the most accurate estimation of the area occupied by cells of each type and not to segment the single individuals. The ‘*segmentation of cells algorithm’* identifies individual cells for subsequent single cell growth estimation (elongation rate). In this case, cells were segmented on the green or the red fluorescent image, according to their fluorescence colour. Single cell location was tracked using an optical flow based algorithm (described below) and the tracking was manually corrected to prevent mistakes. Subparts of the chambers were randomly selected for the single cell segmentation and tracking and 250 cells per chamber were analysed on average, giving a total of 15,475 cells across 61 chambers. The area close (within 8 µm) to the open end of the chamber was not considered for analysis, as amino acid concentrations in this area are lower because they are washed out into the main flow channel. The tracking algorithm based on optical flow can be described in three steps: 1) estimate vector field of movement *M* between subsequent segmented images *S_1_* and *S_2_*using Farneback^47^ algorithm 2) back-transform the second image *S_2,backtransformed_*= - *M* • *S_2_*, to obtain a prediction of how *S_1_* should look like based on the vector field of motion 3) for each cell in *S_1_* determine the area overlap with cells in *S_2,backtransformed_*; cells in *S_1_*are tracked to cells in *S_2_* based on maximum overlap area.

### Cell elongation rate

Cell elongation rates (i.e. growth rates) were calculated by fitting the exponential curve L(t)=L(0) 2^µ·*t*^ to the cell length L over time. The fitting was done using a linear fit on the logarithm of the cell length over a sliding time window of 5 time-points (40 minutes). Length of a cell was measured as the length of the major axis of the ellipse that approximates the cell, i.e. the ellipse that has the same normalized second central moments as the cell.

### Correlation analysis

We quantified the composition of the neighbourhood of a focal cell as the fraction of the partner present in that neighbourhood, e.g. in the case of the tryptophan auxotroph we quantified the fraction ΔproC/(ΔproC+ΔtrpC). ΔtrpC and ΔproC are the areas (in pixel) occupied by each auxotroph, therefore they are a measurement of biomass and not of cell number. To calculate the fraction above, we first identified biomass of the two types as described in the image analysis section; then we calculated the area in pixel that each cell type occupies within increasing distances from the focal cell’s perimeter. For a given distance, we plotted the fraction (x-axis) against the growth rate (y-axis) for all cells and we calculated Spearman’s rank correlation coefficient (no assumption on the functional relationship between variables). The correlation coefficient is maximal at a specific distance, which we call interaction range. We use linear regression to characterize the relationship between the growth rate of the cells and the fraction of the amino acid producing partner present within the estimated interaction range. For Fig. 2d, the correlation is calculated as Spearman ρ on 1,985 data points for proline auxotrophs and 1,769 for tryptophan auxotrophs, both from four biological replicates (with 22 chambers in total). The same analysis performed when cells are fed amino acids shows that growth does not depend on the neighbours when amino acids are present in the medium (Fig. S1).

### Individual-based model

We consider two cell types living on a 40×40 squared grid: the first type can only produce amino acid 1, and its limited in growth by the supply of amino acid 2; the second type can only produce amino acid 2, and its limited in growth by the supply of amino acid 1. We track the spatial distribution of the internal (*I*) and external (*E)* amino acid concentration as function of the location in the monolayer (x, y) and time (t). We expect these concentrations to be constant in the direction perpendicular to the monolayer of cells (z-direction), thus we integrate over the z-direction. For brevity, we will omit the variables (x, y, t) in the notation. We make the following assumptions:

a. Cells maintain a constant internal concentration *I* of the amino acid they can produce.
b. Growth of a cell is limited only by the amino acid the cell cannot produce; growth is modeled using the Monod equation *µ =µ^wt^ I / (I + K),* where *K* is the concentration at which cells grow at half maximum speed.
c. Both cell types can grow at the same rate *µ^wt^* when I>>K, where *µ^wt^* is growth of the wild type. This was experimentally assessed (see Fig. S7).
d. Cells take up amino acids actively^30^, and the process is approximated with linear kinetics: *uptake = r^u^ E*, where *r^u^* is the uptake rate and *E* is the external concentration. Linear kinetics approximates Monod kinetics if the concentrations of external amino acids *E* are low, as is the case in our experimental system.
e. Cells leak amino acids through passive diffusion through the cellular membrane^30^ *leakage = r ^l^ (I - E)*, where *r ^l^* is the leakage rate.
f. Diffusion in the extracellular environment is modeled as diffusion in a crowded environment ^48^ *D*^*eff*^ = *D*(1 − *ρ*)/(1 + *ρ*/*2*), where *D* is the diffusion constant and *ρ* the cell density.
g. The ratio between the volume inside a cell and the available volume outside of a cell is constant and equal to *α* = *ρ* /(1 − *ρ*).

With these assumptions, we can write the following equations for the internal concentration of amino acids for a cell of the first type - which produces amino acid 1, and not amino acid 2:

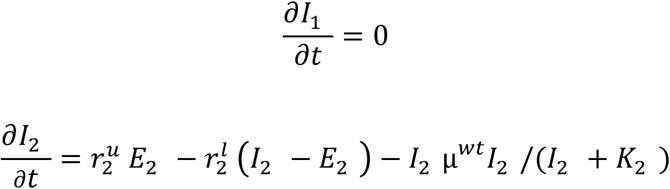

and for the second type - which produces amino acid 2 and not 1:

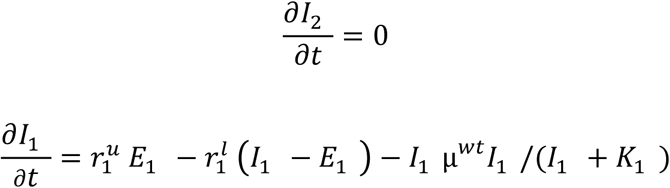

The external concentration of each amino acid is:

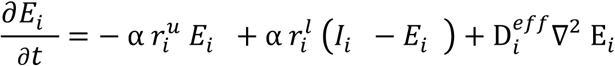

All parameters are taken from literature or are measured, with the exception of the leakage rates, which are estimated from data (see Table S1). These equations can be used to predict cells’ growth rates in real or artificial arrangements of the two cell types. See Supplementary Information, section 3, for a discussion about the effect of these parameters on the length scale of interactions, and for more details on the model.

### Cellular automaton

The cellular automaton models a consortium of two or more types of organisms that live on a grid and benefit from the presence of the other types. The model rests on few general assumptions: first, individuals place offspring close to themselves; second, reproductive success of individuals depends on the fraction of neighbours of the other type within the interaction range, the sole parameter in the model.

An operative description of the cellular automaton follows: individuals reside in a spatially structured setting, each occupying a site on a 40×40 grid; each site has 8 adjacent sites on the grid (Moore neighbourhood) and boundary condition wrap the grid into a torus. For the communities consisting of two types, there are individuals of types 0 and 1. At every time step an individual dies at a random location on the grid and it is replaced with an individual of type 0 or 1. It will be of type 0 with probability P(0):

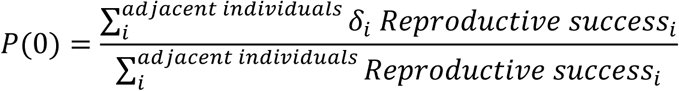

where *δ*_*i*_ is the Dirac delta function, which is one if grid site *i* contains type 0 and zero otherwise. The reproductive success of each individual *i* is:

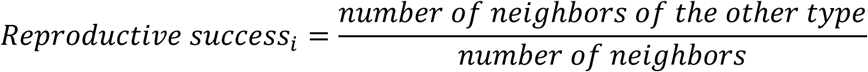

Individuals interact with all other individuals within a neighbourhood of range *R* (a square-shaped neighbourhood). For communities with more than two types, the reproductive success is equal to the fraction of neighbours that is most rare in the neighbourhood:

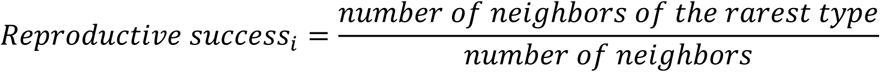

All the rest is easily extended from the two types community described above to communities of more than two types. To compare consortia with a different number of types, the reproductive success is normalized by the reproductive success the consortium has in well-mixed conditions (*R* → ∞), which is 1/2 for two types, 1/3 for three and 1/4 for four.

Starting from different initial configurations and varying proportions of the types, we let the system evolve and stop the simulation after the system has attained a dynamical equilibrium where the average reproductive success of individuals remains approximately constant. Average steady state reproductive success result from 100 independent runs of the cellular automaton. The cellular automaton is implemented in C++.

### Dataset and statistical analysis

The dataset consists of 13,670 cells, from 61 chambers, grouped into ten biological replicates including both fluorescent label combinations. Four biological replicates were done with ΔtrpC-GFP and ΔproC-RFP (consortium1) and six were done with ΔtrpC-RFP and ΔproC-GFP (consortium 2). Each biological replicate corresponds to one channel in a microfluidic chip and for each channel on average 6 chambers were analysed (range: 3-9). Inside each chamber, on average 224 cells were tracked in time, as described in the *Image analysis* method section. The experiments were performed in three independent runs using different microfluidic chips and different batches of media (first chip with four replicates of consortium 1, second chip with two replicates of consortium 2 and third chip with four replicates of consortium 2). The interaction range and relation between growth and neighbourhood were estimated separately for consortium 1 and 2. The interaction ranges are consistent for the two consortia (Fig. 2d shows consortium 1, Fig. S6a shows consortium 2), but the fluorescent label affect the growth rate to some extent: the ΔtrpC-RFP grows generally slower than the ΔtrpC-GFP (Fig. 2f-g shows consortium 1, Fig. S6b-c consortium 2). To assess the variability of the estimate of the interaction range, we repeated the analysis for each replicate in isolation (results are shown in Fig. 2e).

### Mixing and average growth rate in the chambers

The level of mixing of the two cell types in each chamber was measured as the ratio between the length of the boundaries between the two types and the total area they occupy together:

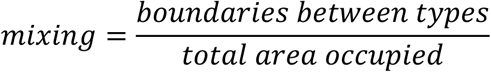

A higher boundaries to area ratio indicates higher levels of mixing. The boundaries between types were estimated using a computationally efficient proxy: we scanned the images in one direction and counted the number of transitions between one type and the other; the total number of transition is a measure of the length of the interface between the two types. In Fig. 5b-c, we normalized the measurement of mixing for our 61 chambers to values between zero (chamber with lowest mixing) and one (chamber with highest mixing).

The average growth rate in the chambers was estimated using a method based on optical flow (using Farneback algorithm^47^). First, a rectangular region was drawn, that had the same width as the chamber and 2/3 of its depth (excluding the third of the chamber close to the opening, where movement of cells is too fast to have a reliable optical flow estimate). As cells grow and flow out of the chamber, they move out of the selected region. Let *B*(*t*) be the biomass in the selected region at time *t*; during a time period Δ*t*, the biomass *B*(*t*) varies due to growth *μ*(*t*) and to flow outside of the selected region Φ(*t*). We can thus write the following equation:

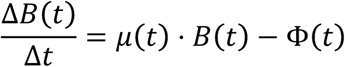

This equation can be used to calculate the growth rate as:

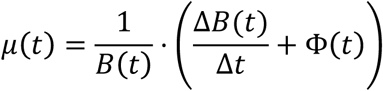

We estimated Φ(*t*) from the two separate fluorescent channels, i.e. we estimated Φ(*t*) as the sum of the optical flow measured on the red and on the green channels separately. All quantities are calculated over a time window Δ*t* = *2h* around time *t = 16h* after the amino acids were removed, and the optical flow is averaged over a strip (20 pixels wide) around the border of the selected region.

## Acknowledgements

We thank Andrea Cavagna for discussing the correlation analysis, Glen D’Souza for advise on the biological assumptions of the model, Gabriele Micali, Roman Stocker, and Michael Doebeli for comments on the model, Kim Schlegel for performing a plate reader experiment, Alex von Wyl for helping correcting cell tracking errors, and Charlotte Brannon for comments on the manuscript. The research was supported by funding from the Swiss National Science foundation (grants no. 31003A_149267 and 31003A_169978 to MA), by an ETH fellowship to DJK, by an Early Postdoc Mobility fellowship from the Swiss National Science Foundation to SVV (grant no. 175123), by ETH Zurich and Eawag.

## Author Contributions

ADC and MA conceived the research, ADC performed the experiment with contributions of SVV, ADC developed the statistical analysis and analysed the data, ADC and DJK developed the image analysis, SVV and ADC conceptualized the individual-based model and SVV implemented it, SS constructed the bacterial strains, DJK constructed the microfluidic device, ADC and MA wrote the manuscript with contributions of SVV.

## Data and materials availability

Source data for all figures is available in the Supplementary Figure Source Data file. A data file containing the full properties of all analysed cells is available on the ETH Research Collection: http://doi.org/10.3929/ethz-b-000367403. Raw image data is available upon request.

## Code availability

The code of the individual based model is available on the Zenodo repository: http://doi.org/10.5281/zenodo.3466038^49^. Additional Matlab scripts for statistical analysis are available upon request.

## References

1. Proulx, S. R., Promislow, D. E. L. & Phillips, P. C. Network thinking in ecology and evolution. Trends Ecol. Evol. 20, 345–353 (2005).

2. Levin, S. A. Ecosystems and the Biosphere as Complex Adaptive Systems Simon. Ecosystems 1, 431–436 (1998).

3. Pickett, S. T. A. & Cadenasso, M. L. Landscape Ecology: Spatial Heterogeneity in Ecological Systems. Science (80-.). 269, 331 LP – 334 (1995).

4. Agrawal, A. A. et al. Filling key gaps in population and community ecology. Front. Ecol. Environ. 5, 145–152 (2007).

5. Falkowski, P. G., Fenchel, T. & Delong, E. F. The Microbial Engines That Drive Earth’s Biogeochemical Cycles. Science (80-.). 320, 1034–1039 (2008).

6. Lynch, S. V & Pedersen, O. The Human Intestinal Microbiome in Health and Disease. N. Engl. J. Med. 375, 2369–2379 (2016).

7. Gore, J. Simple organizing principles in microbial communities. Curr. Opin. Microbiol. 45, 195–202 (2018).

8. Nadell, C. D., Drescher, K. & Foster, K. R. Spatial structure, cooperation and competition in biofilms. Nat. Rev. Microbiol. 14, 589 (2016).

9. Flemming, H.-C. et al. Biofilms: an emergent form of bacterial life. Nat. Rev. Microbiol. 14, 563–575 (2016).

10. Tan, J., Zuniga, C. & Zengler, K. Unraveling interactions in microbial communities - from co-cultures to microbiomes. J. Microbiol. 53, 295–305 (2015).

11. D’Souza, G. et al. Less is more: Selective advantages can explain the prevalent loss of biosynthetic genes in bacteria. Evolution (N. Y). 68, 2559–2570 (2014).

12. Mee, M. T., Collins, J. J., Church, G. M. & Wang, H. H. Syntrophic exchange in synthetic microbial communities. Proc. Natl. Acad. Sci. U. S. A. 111, E2149–56 (2014).

13. Schink, B. Synergistic interactions in the microbial world. Antonie Van Leeuwenhoek Int. J. Gen. Mol. Microbiol 81, 257–261 (2002).

14. Christensen, B. B., Haagensen, J. A. J. J., Heydorn, A. & Molin, S. Metabolic commensalism and competition in a two-species microbial consortium. Appl. Environ. Microbiol. 68, 2495–2502 (2002).

15. Dal Co, A., Ackermann, M. & van Vliet, S. Metabolic activity affects the response of single cells to a nutrient switch in structured populations. J. R. Soc. Interface 16, 20190182 (2019).

16. Rutherford, S. T. & Bassler, B. L. Bacterial quorum sensing: its role in virulence and possibilities for its control. Cold Spring Harb. Perspect. Med. 2, a012427 (2012).

17. Flemming, H.-C. & Wuertz, S. Bacteria and archaea on Earth and their abundance in biofilms. Nat. Rev. Microbiol. 17, 247–260 (2019).

18. Muller, M. J. I., Neugeboren, B. I., Nelson, D. R. & Murray, A. W. Genetic drift opposes mutualism during spatial population expansion. Proc. Natl. Acad. Sci. 111, 1037–1042 (2014).

19. Darch, S. E. et al. Spatial determinants of quorum signaling in a Pseudomonas aeruginosa infection model. Proc. Natl. Acad. Sci. 115, 201719317 (2018).

20. He, X. et al. Microbial interactions in the anaerobic oxidation of methane: model simulations constrained by process rates and activity patterns. Environ. Microbiol. 21, 631–647 (2019).

21. Drescher, K., Nadell, C. D., Stone, H. A., Wingreen, N. S. & Bassler, B. L. Report Solutions to the Public Goods Dilemma in Bacterial Biofilms. Curr. Biol. 24, 50–55 (2014).

22. McGlynn, S. E., Chadwick, G. L., Kempes, C. P. & Orphan, V. J. Single cell activity reveals direct electron transfer in methanotrophic consortia. Nature 526, 531 (2015).

23. Momeni, B., Waite, A. J. & Shou, W. Spatial self-organization favors heterotypic cooperation over cheating. Elife 2, 1–18 (2013).

24. Stump, S. M., Johnson, E. C., Sun, Z. & Klausmeier, C. A. How spatial structure and neighbor uncertainty promote mutualists and weaken black queen effects. J. Theor. Biol. 446, 33–60 (2018).

25. Stump, S. M., Johnson, E. C. & Klausmeier, C. A. Local interactions and self-organized spatial patterns stabilize microbial cross-feeding against cheaters. J. R. Soc. Interface 15, 20170822 (2018).

26. Nowak, M. A., Tarnita, C. E. & Antal, T. Evolutionary dynamics in structured populations. Philos. Trans. R. Soc. B Biol. Sci. 365, 19–30 (2010).

27. Egland, P. G., Palmer, R. J. & Kolenbrander, P. E. Interspecies communication in Streptococcus gordonii-Veillonella atypica biofilms: Signaling in flow conditions requires juxtaposition. Proc. Natl. Acad. Sci. 101, 16917–16922 (2004).

28. Morris, J. J. Black Queen evolution : the role of leakiness in structuring microbial communities. Trends Genet. 31, 475–482 (2015).

29. Oliveira, N. M., Niehus, R. & Foster, K. R. Evolutionary limits to cooperation in microbial communities. PNAS 111, 17941–17946 (2014).

30. D’Souza, G. et al. Ecology and evolution of metabolic cross-feeding interactions in bacteria. Nat. Prod. Rep. 35, 455–488 (2018).

31. Marchal, M. et al. A passive mutualistic interaction promotes the evolution of spatial structure within microbial populations. BMC Evol. Biol. 17, 106 (2017).

32. Dobay, A., Bagheri, H. C., Messina, A., Kümmerli, R. & Rankin, D. J. Interaction effects of cell diffusion, cell density and public goods properties on the evolution of cooperation in digital microbes. J. Evol. Biol. 27, 1869–1877 (2014).

33. Lindsay, R. J., Pawlowska, B. J. & Gudelj, I. When increasing population density can promote the evolution of metabolic cooperation. ISME J. 12, 849– 859 (2018).

34. Ross-gillespie, A. & Kümmerli, R. Collective decision-making in microbes. 5, 1–12 (2014).

35. Stacy, A. et al. Bacterial fight-and-flight responses enhance virulence in a polymicrobial infection. Proc. Natl. Acad. Sci. 111, 7819–7824 (2014).

36. Hol, F. J. H. et al. Spatial Structure Facilitates Cooperation in a Social Dilemma : Empirical Evidence from a Bacterial Community. 8, 2–11 (2013).

37. Mitri, S., Xavier, J. B. & Foster, K. R. Social evolution in multispecies bio films. PNAS 108, 10839–10846 (2011).

38. Davies, D. G. & Geesey, G. G. Regulation of the alginate biosynthesis gene algC in Pseudomonas aeruginosa during biofilm development in continuous culture. Appl. Environ. Microbiol. 61, 860–867 (1995).

39. Saha, M. et al. Microbial siderophores and their potential applications: a review. Environ. Sci. Pollut. Res. 23, 3984–3999 (2016).

40. Leventhal, G. E., Ackermann, M. & Schiessl, K. T. Why microbes secrete molecules to modify their environment: The case of iron-chelating siderophores. J. R. Soc. Interface 16, (2019).

41. DeMalach, N., Zaady, E., Weiner, J. & Kadmon, R. Size asymmetry of resource competition and the structure of plant communities. J. Ecol. 104, 899– 910 (2016).

42. Soliveres, S., Smit, C. & Maestre, F. T. Moving forward on facilitation research: response to changing environments and effects on the diversity, functioning and evolution of plant communities. Biol. Rev. 90, 297–313 (2015).

43. Schmitz, O. J., Miller, J. R. B., Trainor, A. M. & Abrahms, B. Toward a community ecology of landscapes: predicting multiple predator–prey interactions across geographic space. Ecology 98, 2281–2292 (2017).

44. Baba, T. et al. Construction of Escherichia coli K-12 in-frame, single-gene knockout mutants: The Keio collection. Mol. Syst. Biol. 2, (2006).

45. Tomasek, K., Bergmiller, T. & Guet, C. C. Lack of cations in flow cytometry buffers affect fluorescence signals by reducing membrane stability and viability of Escherichia coli strains. J. Biotechnol. 268, 40–52 (2018).

46. Datsenko, K. A. & Wanner, B. L. One-step inactivation of chromosomal genes in Escherichia coli K-12 using PCR products. Proc. Natl. Acad. Sci. 97, 6640– 6645 (2000).

47. Farneback, G. Two-frame motion estimation based on polynomial expansion. Image Anal. Proc. 2749, 363–370 (2003).

48. Lebenhaft, J. R. & Kapral, R. Diffusion-controlled processes among partially absorbing stationary sinks. J. Stat. Phys. 20, 25–56 (1979).

49. Dal Co, A., van Vliet, S., Kiviet, D. J., Schlegel, S. & Ackermann, M. Code for: Short-range interactions govern the dynamics and functions of microbial communities. (2019). doi:10.5281/ZENODO.3466038

