## Supplementary Information for "Short-range interactions govern the dynamics and functions of microbial communities"

**5 Supplementary Tables** **16**

6 Supplementary Video - Caption 16

7 References 17

### 1. Supplementary Discussion

**1.1. Independence of growth rate from neighbours when amino acids are supplied.** We verified that growth of auxotrophic cells does not depend on the identity of their neighbours when amino acids are supplied with the media (Fig. S1). Specifically, we calculated the correlation between growth rate of single cells and the fraction of neighbours of the other type within neighbourhoods of different sizes. We found that the correlation is low for all neighbourhoods analysed. This indicates that the growth rate of auxotrophic cells is not affected by the presence of the other cell type in their surrounding if amino acids are supplied with the media.

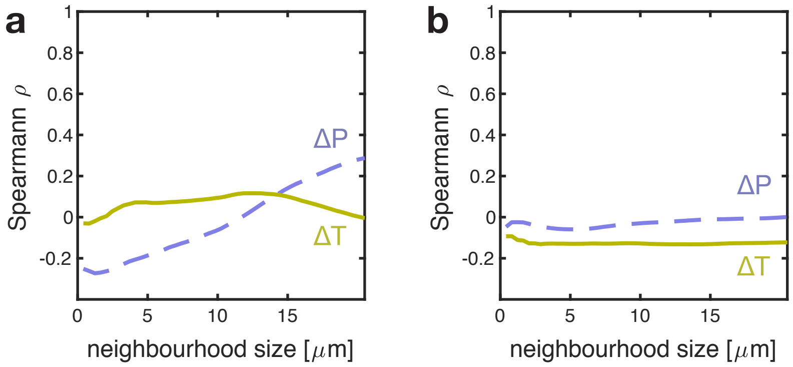

**Fig. S1. Growth does not depend on the identity of neighbours when amino acids are fed.** When media is supplemented with proline and tryptophan, the growth of the auxotrophic cells does not depend on the presence of the partner near by. The correlation between growth rate of cells and fraction of the parter is low for all neighbourhood sizes analysed. Panel (a) shows results for consortium 1 ($\Delta trpC$-GFP and $\Delta proC$-RFP), panel (b) for consortium 2 ($\Delta trpC$-RFP and $\Delta proC$-GFP).

**1.2. Independence of growth rate from distance to the nutrient source.** We tested whether the growth rate of the two auxotrophs depends on the distance from the chamber's opening on the feeding channel, where the media with glucose flows, and we found that there is a very weak association between the two quantities (Fig. S2).

**1.3. Robustness of interaction range estimate to spatial arrangement.** The model allowed us to verify that the interaction range we measure does not arise from the spatial arrangements we analyse, but is rather a property of the system. Generally, inside the communities, the two genotypes form patches because two daughter cells tend to remain close in space after division. The two cell types display different typical patch sizes, with the $\Delta trpC$ (the auxotroph that has the smaller interaction range) forming smaller patches. This observation raises the question: is our correlation analysis affected by patch size? We tested whether patch size affects our estimate of the interaction range by generating several synthetic datasets, each with 100 configurations of the two types arranging in

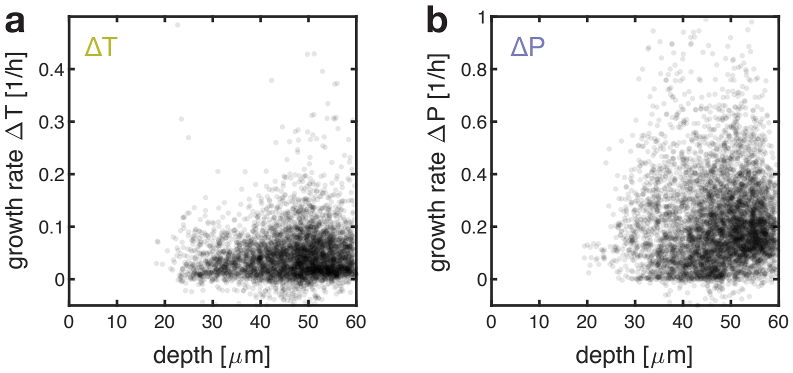

**Fig. S2. Growth of cells does not depend on their distance from the chamber's opening.** The growth of both auxotrophic cells does not correlate with the distance from the chamber's opening into the feeding channel ("depth" in the figure): $\rho = 0.07$,$p < 10^{-4}$, $n = 4567$ for $\Delta trpC$ and $\rho = 0.05$, $p < 10^{-4}$, $n = 5905$ for $\Delta proC$, Spearman.

patches of controlled sizes (Fig. S3a); we analysed these synthetic datasets in the same way as our empirical dataset (Fig. S3b and S3c). The results confirmed that the interaction range of each type, i.e. the location of the correlation peak in Fig. 2d, is robust to changes in patch size (Fig. S3d). In particular, we can show that the location of the peak does not change more than 50% for a range of patch sizes that can be observed in the data (visual inspection). This result supports that our analysis of correlation between growth rates of individuals and their neighbourhood composition is a valid method to determine interaction ranges directly and independently from patch size.

Finally, we also tested if patch size correlates with growth rate of a patch. One could imagine that larger patches imply higher growth rate of the patch. However we verified that patch size does not correlate significantly with growth rate of a patch. We performed a correlation analysis between patch size and growth of a patch for the tryptophan auxotroph (for this cell type it is possible to define closed patches because it is typically in minority in the communities). Larger patches are not found to grow faster, likely because cells inside large patches tend to grow slower (or not at all) as they cannot retrieve the amino acids they need because of the short interaction range.

**1.4. Limitation to the prediction of growth rates.** The model recapitulates quantitatively the effect of spatial arrangement on growth (Fig. 3d - linear correlation between binned growth rates predicted by the model and measured in the data $R^2 > 0.95$), However, the model tends to overestimate the absolute growth rate of cells (in Fig. 3d one line displays an intercept). In fact, the classical Monod equation does not consider that cells may need substrate (the limiting amino acid here) even when they do not grow. For this reason, the original Monod equation is often modified by introducing a term of maintenance[3]. A more refined model including a growth cost could improve the estimation of growth rates; we keep this for future studies. Finally, discrepancies between the estimated and experimentally measured length scales can also originate from the simplified representation of the

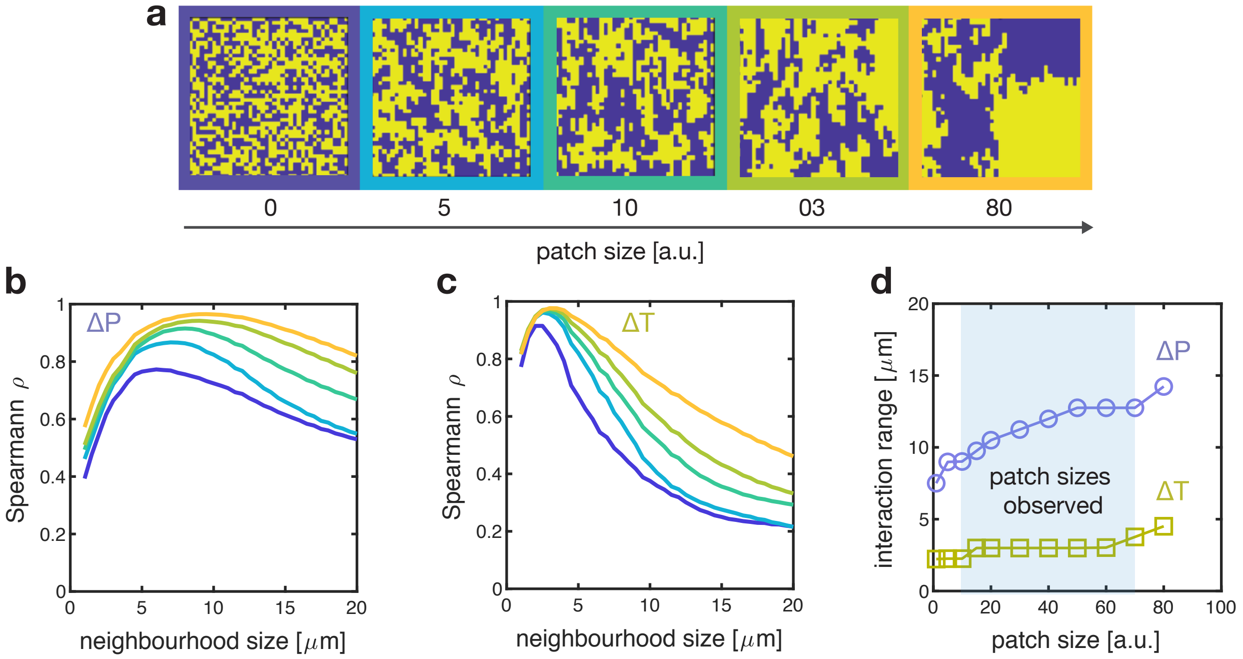

**Fig. S3. Robustness of interaction range estimate to spatial arrangement of types.** (a) Examples of artificial arrangements with controlled patch size; dataset of 100 different arrangements per patch size were generated and analysed. The shape of the correlation curve changes for both proline (b) and tryptophan (c) auxotrophs but the interaction range changes only minimally (d) for a range of patch sizes that can be observed in the data.

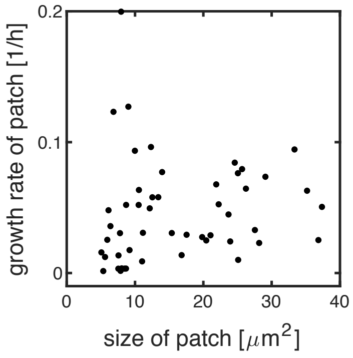

**Fig. S4. Size of a patch does not correlate with its growth rate** The two different genotypes in our system form patches because two daughter cells tend to remain close in space after division. Contrary to intuition, larger patches of the tryptophan auxotrophs do not imply higher growth rate of the patch, because cells in the interior of a large patch tend to grow slower (or not at all) as they cannot retrieve the amino acids they need. $\rho = 0.3$, $p = 0.02$ , $n = 50$, Spearman correlation.

physical space we have in the model (i.e. cells are assumed to be spheres that live on a regularly spaced squared grid).

**1.5. Tradeoff in uptake rates of amino acids.** Our simulations show that increasing the interaction range (by lowering uptake rates) can increase the average growth rate of the two auxotrophs when amino acids do not diffuse out of the system (Fig. 5e). However, when amino acid can diffuse out of the system, there is a tradeoff: when uptake rates are too low, the average growth of the auxotrophs decreases because amino acids tend to diffuse away (Fig. S5). In our chambers, where amino acids can diffuse away, having very low uptake rates and a very large interaction range can reduce growth of cells. Fig. S5 shows that the tryptophan auxotroph (in yellow) grows better if its interaction range increases, while the proline auxotroph (in purple) interacts at range that maximises its growth.

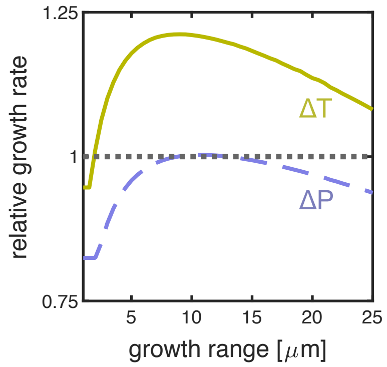

**Fig. S5. Very low uptake rates reduce growth of cells if amino acids can diffuse out of the system.** When simulating systems open on one side, like our chambers, the growth rate of the auxotrophs decreases (relative growth below one) if they have very low uptake rates of the amino acids, i.e. when they have large growth ranges.

While it is intuitive that low uptake rates can decrease growth rate of cells in an open system (where amino acids can diffuse away), it might be unclear why low uptake rates can increase growth rate in a close system (Fig. 5e). For example, one could think that low uptake rates could decrease the internal concentration of amino acids in the cells, thereby decreasing their growth rates. We explain why this is not the case, using analytical arguments (we refer to the notation and the equations in section 3.4). For the symmetric arrangement of the two auxotrophs shown in Fig. 3a (where the interface between the two auxotrophs is a straight line), the growth rate at the interface does not depend on $r^u$ (equation 25). However, the growth rate away from the interface depends on $r^u$. Specifically, we can show that $\epsilon = (r^u + r^l)/r^l * E$ increases at every location when $r^u$ increases. This is because $\epsilon$ at interface between the two types remains the same ($\epsilon = I^c/2$), but $\epsilon$ decays slower away from the interface. Higher $\epsilon$ implies higher concentrations of $I_L$, the amino acids that limits growth rate of cells, because $I_L(\epsilon)$ is an increasing function of $\epsilon$ (and does not depend on

$r^u$, see equation 22). Higher $I_L$ for every cell finally implies higher growth rates $\mu$ for every cell, because $\mu \propto \frac{I_L}{K+I_L}$. We conclude that, in the symmetric arrangement shown in Fig. 3a, lowering $r^u$ increases the growth rate of every cell. A similar analytical argument for any arbitrary arrangement is too complicated, but our simulations show that community growth always increases with lower $r^u$ (Fig. 4c). Overall this suggests that the growth rate of cells is higher when the uptake rate of amino acids $r^u$ is lower, as long as we consider closed systems where amino acids cannot diffuse away.

### 2. Supplementary Data

We checked that the fluorescent marker the strains carried did not affect our main results. Fig. S6 show the correlation analysis between single cells growth rates and fraction of the partner in the interaction range for six biological replicates done with $\Delta trpC$-RFP and $\Delta proC$-GFP (consortium 2); Fig. 2 in the main text shows four biological replicates done with $\Delta trpC$-GFP and $\Delta proC$-RFP (consortium 1). The interaction ranges are consistent for the two consortia (Fig. 2d shows consortium 1, S6a consortium 2), but the fluorescent label affect the growth rate to some extent: the $\Delta trpC$-RFP grows generally slower than the $\Delta trpC$-GFP (Fig. 2f-g shows consortium 1, Fig. S6b-c consortium 2). In batch cultures $\Delta trpC$-RFP also has a slight growth defect: it is the only strain that grows significantly slower than the wild type (Fig. S7). However, the difference in growth rate between $\Delta trpC$-RFP ($\mu$=1.16 1/h) and $\Delta trpC$-GFP ($\mu$=1.21 1/h) in batch cultures is not significant (Fig. S7).

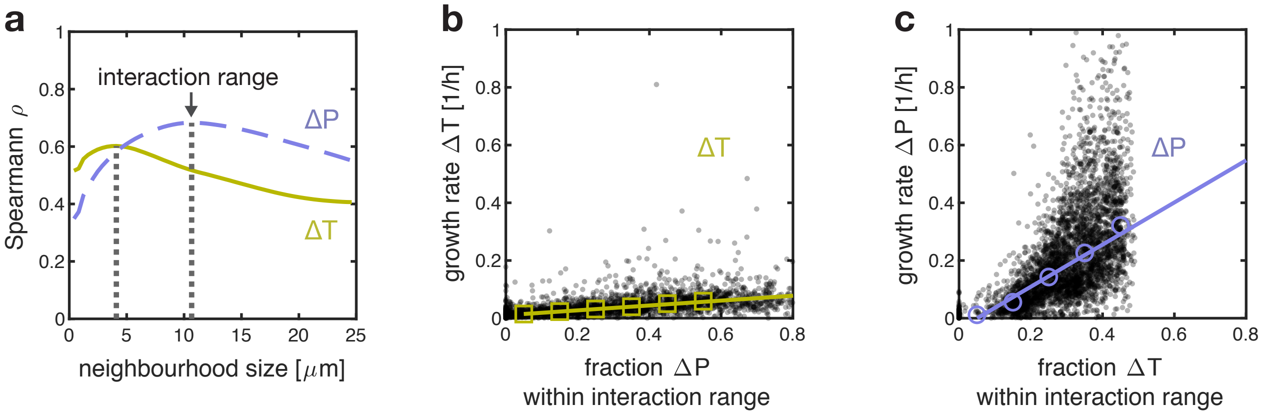

**Fig. S6. Individuals interact at a small spatial range.** All three panels shows data for consortium 2 ($\Delta trpC$-RFP and $\Delta proC$-GFP), and complement Fig. 2 showing data from consortium 1 ($\Delta trpC$-GFP and $\Delta proC$-RFP). (a) The cells' growth rate correlates maximally with the identity of their neighbours within the interaction range. (b-c) Both auxotrophic cells grow faster when surrounded by more complementary partners inside the interaction range. Tryptophan auxotrophs (b) achieve generally smaller growth rates then proline auxotrophs (c), as shown by the slopes of the linear regression (0.79 for $\Delta proC$ and 0.089 for $\Delta trpC$). Black dots: single cells (3,920 for $\Delta proC$ and 2,798 for $\Delta trpC$ ); open symbols: binned median values; lines: linear regression on binned values.

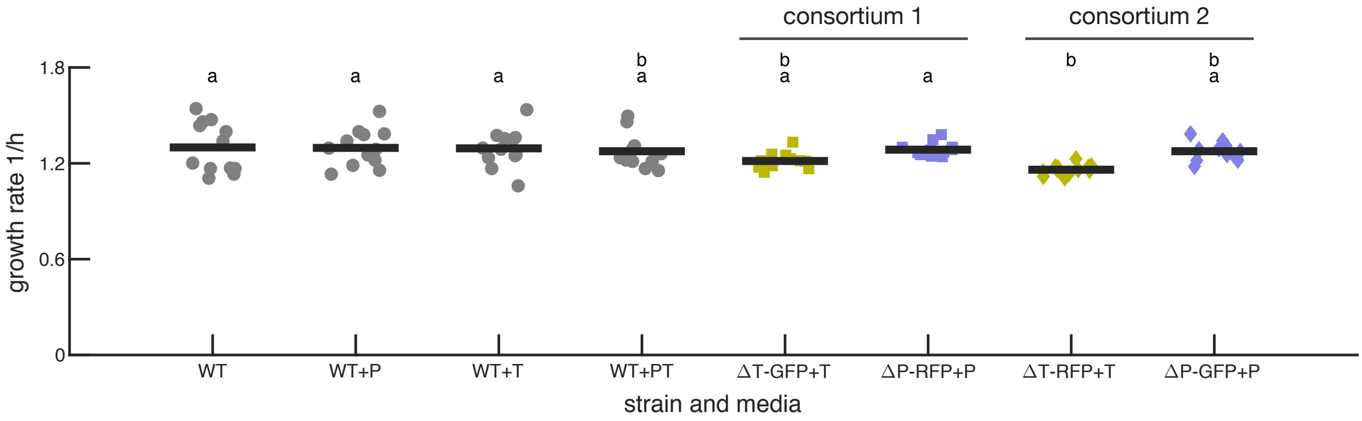

**Fig. S7. Growth rate in batch cultures of wild type and auxotroph strains.** Wild type cells grow at a similar rate in M9 glucose medium without amino acids (WT) and in M9 glucose medium supplemented with 50 $\mu g/mL$ of proline (WT+P), 20 $\mu g/mL$ of tryptophan (WT+T), or both (WT+PT). The auxotrophic mutants in consortium 1 (ΔT-GFP + ΔP-RFP) grow at the same rate as the wild type. The proline auxotroph was grown in medium supplemented with 50 $\mu g/mL$ of proline and the tryptophan auxotroph in medium supplemented with 20 $\mu g/mL$ of tryptophan. Markers show maximal growth rates in batch culture of individual replicates ($n = 12$), bars show average values. Shared letters (top of panel) indicate no significant difference in growth rate (ANOVA analysis with post-hoc Tukey-Kramer pairwise comparison, $F = 3.29$, $df = 7$, $p = 4 \cdot 10^{-3}$). For the wild type there was no significant difference between the strains with the different color labels (TB205 with RFP label and TB204 with GFP label, ANOVA analysis on growth rate data of wild type in all four growth media, testing for the effect of medium [$F = 2.53$, $df = 3$, $p = 0.07$] and color label [$F = 0.02$, $df = 1$, $p = 0.88$]); the data of both strains was thus pooled together.

### 3. Supplementary Equations

**3.1. Rescaled equations for the individual-based model.** The individual-based model describes two cell types which we call here type A and type B. Type A can only produce amino acid 1 while type B can only produce amino acid 2. The growth of type A is thus limited by the supply of amino acid 2 leaked by type B cells and vice versa. We assume that cells grow as a monolayer within a three-dimensional space (i.e. as they do in the growth chambers). We expect that the concentration of amino acids is constant in the direction perpendicular to the monolayer of cells ($z$-direction). We can thus integrate over the $z$-direction and track the spatial distribution of the internal $I$ and external $E$ amino acid concentration as function of the location in the monolayer $(x, y)$ and time $(t)$. We model cells as spherical objects that are arranged on a regularly spaced squared grid. We model the following processes (the subscript $i$ refers to amino acid 1 or 2):

- Cells actively take up amino acids from the external environment following a first order kinetic with uptake rate $r_i^u$: $uptake = r_i^u \cdot E_i$. First order kinetics are a good approximation of Michaelis-Menten kinetics when external concentrations are low: $uptake = \frac{v_{max} \cdot E}{K_{up}+E} \approx \frac{v_{max}}{K_{up}} \cdot E_i + \mathcal{O}(E_i^2) \equiv r_i^u \cdot E_i$.
- Cells leak amino acids passively with leakage rate $r_i^l$: $leakage = r_i^l \cdot (I_i - E_i)$
- Cells consume amino acids proportional to their growth rate: $consumption = \mu \cdot I_i$. Cell growth is limited by the amino acid that they cannot produce, following a Monod kinetic: $\mu = \frac{\mu_i \cdot I_i}{K_i + I_i}$, where $\mu_i$ is the maximum growth rate of cells that cannot produce amino acid $i$.
- Cells produce amino acids. We assume that cells maintain a constant concentration $I_i^C$ of the amino acid that they can produce as a result of tight regulation of the amino acid production rates (most amino acids synthesis pathways are regulated using end-product inhibition[1,2]).
- Amino acids diffuse through the external environment with the effective diffusion constant $D_i^{eff} = \frac{(1-\rho) \cdot D_i}{(1+\frac{\rho}{2})}$, where $D_i$ is the diffusion constant in empty space and $\rho$ the cell density.

Using these assumptions we can write down the dynamical equations for the internal $I_i(x, y, t)$ and external $E_i(x, y, t)$ amino acid concentration:

$$\frac{\partial I_1}{\partial t} = 0 \qquad \text{if type is A} \qquad [1]$$

$$\frac{\partial I_1}{\partial t} = r_1^u \cdot E_1 - r_1^l \cdot (I_1 - E_1) - \frac{\mu_1 \cdot I_1}{K_1 + I_1} \cdot I_1 \qquad \text{if type is B} \qquad [2]$$

$$\frac{\partial I_2}{\partial t} = r_2^u \cdot E_2 - r_2^l \cdot (I_2 - E_2) - \frac{\mu_2 \cdot I_2}{K_2 + I_2} \cdot I_2 \qquad \text{if type is A} \qquad [3]$$

$$\frac{\partial I_2}{\partial t} = 0 \qquad \text{if type is B} \qquad [4]$$

$$\frac{\partial E_i}{\partial t} = -\alpha \cdot r_i^u \cdot E_i + \alpha \cdot r_i^l \cdot (I_i - E_i) + D_i^{eff} \nabla^2 E_i \qquad [5]$$

Where $\alpha = \frac{V_{in}}{V_{out}} = \frac{\rho}{1-\rho}$ is the ratio between the intra and extra cellular volume. We can reduce the number of parameters of our model by measuring all amino acid concentrations relative to their Monod constant: $\mathcal{I}_i = \frac{I_i}{K_i}$ and $\mathcal{E}_i = \frac{E_i}{K_i}$. This gives the following equations:

$$\frac{\partial \mathcal{I}_1}{\partial t} = 0 \qquad \text{if type is A} \qquad [6]$$

$$\frac{\partial \mathcal{I}_1}{\partial t} = r_1^u \cdot \mathcal{E}_1 - r_1^l \cdot (\mathcal{I}_1 - \mathcal{E}_1) - \frac{\mu_1 \cdot \mathcal{I}_1}{1 + \mathcal{I}_1} \cdot \mathcal{I}_1 \qquad \text{if type is B} \qquad [7]$$

$$\frac{\partial \mathcal{I}_2}{\partial t} = r_2^u \cdot \mathcal{E}_2 - r_2^l \cdot (\mathcal{I}_2 - \mathcal{E}_2) - \frac{\mu_2 \cdot \mathcal{I}_2}{1 + \mathcal{I}_2} \cdot \mathcal{I}_2 \qquad \text{if type is A} \qquad [8]$$

$$\frac{\partial \mathcal{I}_2}{\partial t} = 0 \qquad \text{if type is B} \qquad [9]$$

$$\frac{\partial \mathcal{E}_i}{\partial t} = -\alpha \cdot r_i^u \cdot \mathcal{E}_i + \alpha \cdot r_i^l \cdot (\mathcal{I}_i - \mathcal{E}_i) + D_i^{eff} \nabla^2 \mathcal{E}_i \qquad [10]$$

In our simulations we further always assume that both auxotrophs have the same maximum growth rate: $\mu_2 = \mu_1 = \mu^{wt}$. This assumption is supported by the experimental observation that both auxotrophs have similar growth rates to each other and to the wild type in batch cultures (Fig. S7). To maintain generality, we will not use this assumption in the remainder of this text.

**3.2. Steady state equations.** We are primarily interested in finding how the growth rate of a cell depends on the arrangement of the different cell types in space. To do this, we have to solve for the steady state distribution of the external $\mathcal{E}_i(x, y)$ and internal $\mathcal{I}_i(x, y)$ amino acid concentration. From the spatial distribution of the internal amino acid concentration we can then calculate the growth rate for each cell and investigate how it depends on the spatial arrangement of the cell types.

We first solve for the steady state concentration of the growth limiting amino by setting equation 7 or 8 to zero:

$$0 = r_i^u \cdot \mathcal{E}_i - r_i^l \cdot (\mathcal{I}_i - \mathcal{E}_i) - \frac{\mu_i \cdot \mathcal{I}_i}{1 + \mathcal{I}_i} \cdot \mathcal{I}_i$$

Which we can solve to find the internal concentration $\mathcal{I}_i^{lim}(\mathcal{E}_i)$ of the limiting amino acid (i.e. the amino acid that a cell cannot produce) as function of the external concentration of that amino acid:

$$\mathcal{I}_i^{lim}(\mathcal{E}_i) = \frac{(r_i^u + r_i^l)\mathcal{E}_i - r_i^l + \sqrt{\left((r_i^u + r_i^l)\mathcal{E}_i + r_i^l\right)^2 + 4(r_i^u + r_i^l)\mu_i\mathcal{E}_i}}{2(\mu_i + r_i^l)} \qquad [11]$$

Next we solve for the steady state concentration of the produced amino acid $\mathcal{I}_i^{prod}(\mathcal{E}_i)$ by solving for the steady state of equation 6 or 9:

$$\mathcal{I}_i^{prod}(\mathcal{E}_i) = \mathcal{I}_i^C \qquad [12]$$

where $\mathcal{I}^C = I^C/K_i$ is the constant internal concentration of the produced amino acid, relative to the Monod constant of that amino acid. Finally we solve for the steady state distribution of the external concentration by setting equation 10 to steady state:

$$\nabla^2\mathcal{E}_i = \frac{\alpha \cdot (r_i^u + r_i^l)}{D_i^{eff}} \cdot \mathcal{E}_i - \frac{\alpha \cdot r_i^l}{D_i^{eff}} \cdot \mathcal{I}_i(\mathcal{E}_i) \qquad [13]$$

where $\mathcal{I}_i(\mathcal{E}_i)$ is given by $\mathcal{I}_i^{prod}(\mathcal{E}_i)$ (equation 12) for grid sites where amino acid $i$ is produced and by $\mathcal{I}_i^{lim}(\mathcal{E}_i)$ (equation 11) otherwise.

We describe the spatial arrangement of the two cell types with the function $T(x,y)$:

$$\begin{aligned} T(x,y) &= 0 \qquad \text{if site } (x,y) \text{ is occupied by type A} \\ T(x,y) &= 1 \qquad \text{if site } (x,y) \text{ is occupied by type B} \end{aligned} \qquad [14]$$

Using this notation we can rewrite equation 13 by explicitly specifying $\mathcal{I}_i(\mathcal{E}_i)$. We then find that the external concentration of the two amino acid is the solution of the equations:

$$\nabla^2\mathcal{E}_1(x,y) = \frac{\alpha(r_1^u + r_1^l)}{D_1^{eff}} \cdot \mathcal{E}_1(x,y) - \frac{\alpha r_1^l}{D_1^{eff}} \cdot \left(T(x,y) \cdot \mathcal{I}_1^{lim}\left(\mathcal{E}_1(x,y)\right) + [1 - T(x,y)] \cdot \mathcal{I}_1^C\right) \qquad [15]$$

$$\nabla^2\mathcal{E}_2(x,y) = \frac{\alpha(r_2^u + r_2^l)}{D_2^{eff}} \cdot \mathcal{E}_2(x,y) - \frac{\alpha r_2^l}{D_2^{eff}} \cdot \left([1 - T(x,y)] \cdot \mathcal{I}_2^{lim}\left(\mathcal{E}_2(x,y)\right) + T(x,y) \cdot \mathcal{I}_2^C\right) \qquad [16]$$

After solving for $\mathcal{E}_i(x,y)$ we can obtain the growth profile $\mu(x,y)$:

$$\mu(x,y) = [1 - T(x,y)] \cdot \frac{\mu_2 \cdot \mathcal{I}_2^{lim}(x,y)}{1 + \mathcal{I}_2^{lim}(x,y)} + T(x,y) \cdot \frac{\mu_1 \cdot \mathcal{I}_1^{lim}(x,y)}{1 + \mathcal{I}_1^{lim}(x,y)} \qquad [17]$$

where $\mathcal{I}_i^{lim}(x,y) = \mathcal{I}_i^{lim}(\mathcal{E}_i(x,y))$ is given by equation 11.

**3.3. Numerical solution and boundary condition.** We numerically solved equations 15 and 16 for an environment that closely matches the experimental growth chambers. Cells were placed on square grid of 40x40 sides for a total of 1600 cells; this number is comparable to the total number of cells we observed in the experimental growth chambers. On one edge of the grid we implement a Dirichlet boundary condition and set $E_i = 0$ to represent the flow-channel where all excreted amino-acids are washed away; on all other edges we implement Neumann no-flux boundary conditions to represents the solid wall of the growth chamber. We discretized equations 15 and 16 using a second order finite difference scheme and solved them using a successive over-relaxation solver. To ensure numerical stability, we implemented a grid-refinement procedure: we first solved the equations on the 40x40 grid and then we iterated on refined grids (successively doubling the number of grid points in each dimension); we used the solution of the previous iteration as the initial state for the successive refined grid. The solution on the refined grid was downsampled to the 40x40 grid to calculate the growth rate for each cell using equation 17 and we continued this procedure until the maximum per cell change in growth rate was less than 1%. All code was implemented in Matlab.

**3.4. Analytical limits.** We found several analytical approximations to our model that give insight into how cell growth and interaction range depend on the model parameters. In this section we derive the approximations, a summary of the main findings is given in section 3.5.

For the derivation of the analytical approximations, we will consider a single cell type at a time and follow only the internal, $I_L$, and external, $E_L$, concentration of the limiting amino-acid for this cell type:

$$\frac{\partial \mathcal{I}_L}{\partial t} = r^u \cdot \mathcal{E}_L - r^l \cdot (\mathcal{I}_L - \mathcal{E}_L) - \frac{\mu^{aux} \cdot \mathcal{I}_L}{1 + \mathcal{I}_L} \cdot \mathcal{I}_L \quad [18]$$

$$\frac{\partial \mathcal{E}_L}{\partial t} = -\alpha \cdot r^u \cdot \mathcal{E}_L + \alpha \cdot r^l \cdot (\mathcal{I}_L - \mathcal{E}_L) + D^{eff} \nabla^2 \mathcal{E}_L. \quad [19]$$

Where $r^u$, $r^l$, and $D^{eff}$ always refer to the uptake, leakage, and effective diffusion constant of the growth limiting amino acid and where $\mu^{aux}$ is the growth rate of a cell that is auxotrophic for this amino acid. We can simplify the notation by rewriting these equation in terms of $\epsilon \equiv \frac{r^u + r^l}{r^l} \mathcal{E}_L$:

$$\frac{\partial \mathcal{I}_L}{\partial t} = r^l \cdot (\epsilon - \mathcal{I}_L) - \frac{\mu^{aux} \cdot \mathcal{I}_L}{1 + \mathcal{I}_L} \cdot \mathcal{I}_L \quad [20]$$

$$\frac{\partial \epsilon}{\partial t} = -\alpha r^l \cdot (\epsilon - \mathcal{I}_L) + D^{eff} \nabla^2 \epsilon \quad [21]$$

Setting the time derivatives to zero and solving 20 for $\mathcal{I}_L$ gives:

$$\mathcal{I}_L(\epsilon) = \frac{r^l(\epsilon - 1) + \sqrt{(r^l)^2(\epsilon + 1)^2 + 4\mu^{aux} r^l \epsilon}}{2\left(\mu^{aux} + r^l\right)} \quad [22]$$

***3.4.1. Maximum cell growth rate.*** Here we derive the analytical expression for the growth rate of an auxotrophic cells surrounded by a large number of producing partners. If we assume that the single auxotroph has a negligible influence on the external concentration (i.e. all space is occupied by producers which have $\mathcal{I} = \mathcal{I}^C$), equation 21 gives the steady state external concentration of amino acids:

$$\epsilon_{max} = \mathcal{I}^C \qquad [23]$$

substituting $\epsilon_{max}$ for $\epsilon$ in eq. 22 we find:

$$\begin{aligned} \mathcal{I}_L^{max} &= \frac{r^l(\mathcal{I}^C - 1) + \sqrt{(r^l)^2(\mathcal{I}^C + 1)^2 + 4\mu^{aux} r^l \mathcal{I}^C}}{2\left(\mu^{aux} + r^l\right)} \\ \mu^{max} &= \frac{\mathcal{I}_L^{max}}{1 + \mathcal{I}_L^{max}} \end{aligned} \qquad [24]$$

This is the growth rate of a single auxotrophs surrounded by a large number of amino acid producing partners. We can simplify this expression if we make the following two assumptions:

**Assumption 1** $\mathcal{I}^C \gg 1$. *Biologically this means that a wild type (amino acid producing) cell can grow nearly as fast in the absence of amino acids ($\mu = \frac{\mu^{wt} \cdot \mathcal{I}^C}{1+\mathcal{I}^C}$) as in the presence of amino acids ($\mu = \mu^{wt}$). We verified experimentally that this assumption holds (Fig. S7).*

**Assumption 2** $r^l \ll \mu^{aux}$. *Biologically this means that in wild type cells the decrease in the concentration of limiting amino acid due to leakage (with rate $r^l$) is small compared to the decrease due to growth (with rate $\mu^{wt} \approx \mu^{aux}$).*

With these assumptions the growth rate of an auxotroph surrounded by the producing partner (eq. 24) simplifies to:

$$\mu^{max} \approx \frac{r^l \mathcal{I}^C}{2} \left( \sqrt{1 + \frac{4\mu^{aux}}{r^l \mathcal{I}^C}} - 1 \right) \qquad [25]$$

***3.4.2. Estimating leakage rates.*** In this subsection we show how we estimated the leakage rates from the maximum empirical growth rates of the auxotrophs. In the limit of $r^l \ll \mu^{aux}$ and $\mathcal{I}^C \gg 1$, the maximum growth rate (equations 25) depends on the product $\mathcal{I}^C \cdot r^l$, and not on the two parameters separately. Both parameters are unknown but we expect $\mathcal{I}^C \gg 1$ (assumption 1), and we arbitrarily set $\mathcal{I}^C = 20$ for both amino acids. This is equivalent to stating that a wild type cell grown in the absence of amino acids can grow at 95% of the growth rate of a wild type cell grown in the presence of amino acids. In experiments we could not find a significant difference in the wild-type growth rate grown with or without amino acids, verifying that this is a reasonable assumption (Fig. S7).

With this, we can estimate the leakage rates $r^l$ for each amino acid from $\mu^{max}$ of the corresponding auxotroph. Note that our result are robust to changes in the value assigned to $\mathcal{I}^C$ as long as it is larger than one (we confirmed that our simulations depend only on the product $\mathcal{I}^C \cdot r^l$ as long as $r^l \ll \mu^{aux}$ and $\mathcal{I}^C \gg 1$).

The maximum empirical growth rate $\mu^{max}$ is estimated for each auxotroph by performing a linear regression between the auxotroph's growth rate and the fraction of the producing partner within the interaction range (Fig. 2f-g); the maximum growth rate is the value extrapolated when the fraction is equal to one. We use linear regression because it is less sensitive to measurement noise than using the maximal observed growth rate, and because very few cells are found completely surrounded by the producing partner because of kin clustering. We verified that the interaction range does not vary substantially in a large range of leakage rates (Fig. S8).

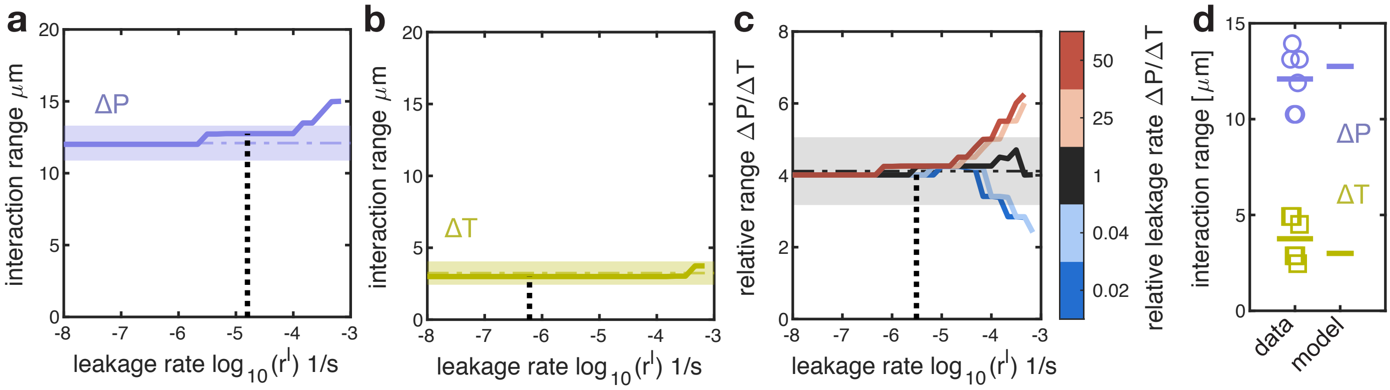

**Fig. S8. The interaction range varies minimally with the leakage rate.** (a-b) The predicted interaction range is consistent with the experimentally measured one across a large range of possible leakage rates. The interaction range of the two auxotrophs was predicted using the model while varying the leakage rates of the amino acids. The model was solved on experimentally measured spatial arrangements. The dashed horizontal lines and shaded regions indicate the mean and 95% confidence interval of the experimentally measured interaction ranges. The dashed vertical line indicates the fitted leakage rate used in all simulations. (c) The predicted relative interaction range is consistent with the experimentally measured one over a large range of possible leakage rates. The relative interaction range (interaction range of $\Delta P$ divided by that of $\Delta T$) was predicted using the model while varying the leakage rates of the amino acids. The $x$-axis shows the geometric mean value of the leakage rate $r^l = \sqrt{r^l_{\Delta P} \cdot r^l_{\Delta T}}$, the different colored lines show different ratios of the leakage rates of the two amino acids $\frac{r^l_{\Delta P}}{r^l_{\Delta T}}$. The dashed horizontal line and shaded region indicate the mean and 95% confidence interval of the experimentally measured relative range. The dashed vertical line indicates the fitted leakage rate used in all simulations (fitted $\frac{r^l_{\Delta P}}{r^l_{\Delta T}} = 26$). (d) Cross-validation of model. The leakage rate was fitted to data from consortium 1 and the interaction range was calculated using the model. This prediction (based solely on data from consortium 1) was compared to the experimentally measured interaction range in consortium 2. The model can quantitatively predict the interaction range ($p = 0.16, n = 6$ for $\Delta T$ and $p = 0.35, n = 6$ for $\Delta P$, t-test)

***3.4.3. Analytical expression for the growth range.*** In this subsection we derive an analytical expression for the length scale over which cells can interact. For most spatial arrangements our model is too complex to be solved analytically. However we can find an approximate analytical solution for the symmetric spatial arrangement shown in Fig. 4a, where the two cell types are segregated in space and meet at a straight interface. The symmetry of this arrangement reduces the problem to one dimension and allows us to find an approximate solution for the amino acid profiles in space. From this we can calculate how the growth rate of cells decrease with the distance from the interface and we define the *growth range* as the length scale over which growth rates decrease by 50%. We confirmed that this growth range (calculated analytically for the symmetric spatial arrangement) is proportional to the interaction range (calculated numerically for complex spatial arrangements, Fig. 4b). The analytical expression for the growth range can thus inform is how the interaction range depends on the model parameters.

We consider the arrangement of cells as shown in Fig. 4a, where the identity of cell types is constant with $y$. This reduces the problem to one-dimension, $x$, which measures the distance of a cell to the interface. We will derive an analytical approximation for the growth profile of cells to the right of the interface ($x > 0$). These cells are growth limited by the amino acid produced by cells to the left of the interface ($x < 0$). We only consider the amino acid that limits growth for cells at $x > 0$, its profile can be found by solving for the steady state of equation 21:

$$D^{eff}\frac{d^2\epsilon(x)}{dx^2} = \alpha r^l \cdot (\epsilon(x) - \mathcal{I}(x)) \qquad [26]$$

Cells at $x < 0$ can produce this amino acid, and their internal concentration is thus given by $\mathcal{I}(x) = \mathcal{I}^C$. Cells at $x > 0$ cannot produce this amino acid, and their internal concentration is thus determined by the external concentration: $\mathcal{I}(x) = \mathcal{I}_L(\epsilon(x))$, where $\mathcal{I}_L(\epsilon)$ is given by 22. We thus have to solve the following equation:

$$\frac{d^2\epsilon}{dx^2} = \begin{cases} \frac{1}{r_0^2}\left(\epsilon - \mathcal{I}^C\right) & \text{if } x < 0 \\ \frac{1}{r_0^2}\left(\epsilon - \mathcal{I}_L(\epsilon)\right) & \text{if } x > 0 \end{cases} \qquad [27]$$

where

$$r_0 = \sqrt{\frac{D^{eff}}{\alpha(r^u + r^l)}}$$

For $x > 0$ the analytical solution of equation 27 cannot be found due to the non-linear term $\mathcal{I}_L(\epsilon)$ (given by equation 22). However it is easy to show that

$$\mathcal{I}_L(\epsilon) < \frac{r^l\epsilon + \mu^{aux}}{r^l + \mu^{aux}}$$

The non linear term $\mathcal{I}_L(\epsilon)$ is thus negligible compared to $\epsilon$ when $\epsilon \gg 1$. For $x \ll 0$ the external concentration is the steady state concentration as found in a region of producers only, i.e. $\epsilon(x \ll 0) = \mathcal{I}^C$ (see subsection 3.4.1), while for $x \gg 0$ all amino acids will have been consumed and

$\epsilon(x \gg 0) = 0$. Because of symmetry we thus expect that at the interface $\epsilon(x=0) \approx \frac{1}{2}\mathcal{I}^C$. As we assumed that $\mathcal{I}^C \gg 1$ (assumption 1) we can thus ignore the non-linear term in equation 27 close to the interface and instead solve the simplified linear ODE:

$$\frac{d^2\epsilon}{dx^2} = \begin{cases} \frac{1}{r_0^2} \cdot \left(\epsilon - \mathcal{I}^C\right) & \text{if } x < 0 \\ \frac{1}{r_0^2} \cdot \epsilon & \text{if } x > 0 \end{cases} \tag{28}$$

These equation can be solved analytically to find:

$$\epsilon(x) = \begin{cases} C_1 \cdot e^{x/r_0} + \mathcal{I}^C & \text{if } x < 0 \\ C_2 \cdot e^{-x/r_0} & \text{if } x > 0. \end{cases}$$

We can solve for $C_1$ and $C_2$ by imposing continuity of concentration and flux at the interface:

$$C_1 \cdot e^{x/r_0} + \mathcal{I}^C|_{x=0} = C_2 \cdot e^{-x/r_0}|_{x=0}$$
$$\frac{C_1}{r_0} \cdot e^{x/r_0}|_{x=0} = -\frac{C_2}{r_0} \cdot e^{-x/r_0}|_{x=0}.$$

From which we find that $C_1 = -\frac{\mathcal{I}^C}{2}$ and $C_2 = \frac{\mathcal{I}^C}{2}$. The external concentration is given thus by:

$$\epsilon(x) = \begin{cases} \mathcal{I}^C \left(1 - \frac{1}{2} \cdot e^{x/r_0}\right) & \text{if } x < 0 \\ \frac{\mathcal{I}^C}{2} \cdot e^{-x/r_0} & \text{if } x > 0 \end{cases} \tag{29}$$

Within the consumer region the amino acid concentration ($\mathcal{E} = \frac{r^l}{r^u + r^l}\epsilon$) decreases exponentially with scale factor $r_0$.

We are now interested in finding an analytical approximation for the *growth range* ($GR$), which is the distance from the interface where cells have 50% of the growth rate they have at the interface:

$$\mu(x = GR) = \frac{1}{2} \cdot \mu(x = 0) \tag{30}$$

as $\mu = \frac{\mu^{aux} \cdot \mathcal{I}}{1+\mathcal{I}}$ it follows that

$$\mathcal{I}_L\Big|_{x=GR} = \frac{\mathcal{I}_L\Big|_{x=0}}{2 + \mathcal{I}_L\Big|_{x=0}} \tag{31}$$

where $\mathcal{I}_L\Big|_x \equiv \mathcal{I}_L(\epsilon(x))$. Substituting equation 22 for $\mathcal{I}_L(\epsilon)$ and equation 29 for $\epsilon(x)$ we can thus solve equation 31 for the growth range:

$$GR = r_0 \cdot \ln\left[\frac{(\mathcal{I}^C - 4)\left(r^l(\mathcal{I}^C + 2) + \sqrt{(\mathcal{I}^C + 2)^2(r^l)^2 + 8\mathcal{I}^C r^l \mu^{aux}}\right) + 8\mathcal{I}^C(r^l + \mu^{aux})}{2(\mathcal{I}^C(\mu^{aux} + 2r^l) - 4r^l)}\right] \tag{32}$$

using assumptions 1 ($\mathcal{I}^C \gg 1$) and 2 ($r^l \ll \mu^{aux}$) this can be simplified to:

$$GR \approx \sqrt{\frac{D^{eff}}{\alpha(r^u + r^l)}} \cdot \ln\left[\frac{r^l \mathcal{I}^C}{2\mu^{aux}}\left(1 + \sqrt{1 + \frac{8\mu^{aux}}{r^l \mathcal{I}^C}}\right) + 4\right] \quad [33]$$

Fig. S9b shows the error of this analytical approximation (eq. 33) of the growth range. As long as the growth range is smaller than 20, our simulations match the analytical result very well. As the growth range approaches 20 (Fig. S9a), the relative error increases because of the finite size (40x40) of the chamber in our simulations; more specifically, the no-flux boundary conditions lead to overestimation of the growth range in the simulations compared to the analytical model.

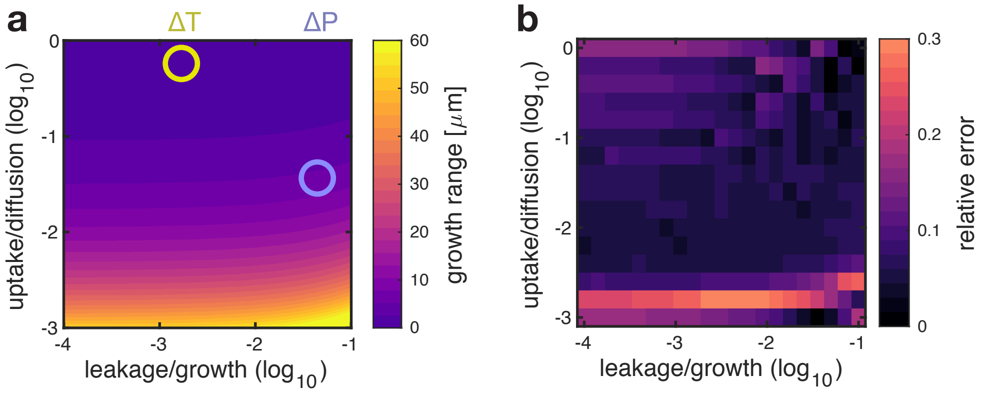

**Fig. S9. Analytical approximation of growth range and simulations agree.** The heat map (b) shows the relative error between analytical approximation (eq. 33) for the growth range and the growth range estimated with simulations. The heat map (a) shows the analytical estimate of the growth range, and shows that the relative error in (b) is low when the growth range is below 20. Purple circle is proline auxotroph and yellow circle is tryptophan auxotroph. Panel a shows the same data as Fig. 4c.

**3.5. Discussion on the effect of parameters.** We have found an analytical approximations for the maximum growth rate of each auxotroph when surrounded by a large number of the amino acid producing partner and for the growth range:

$$\mu^{max} \approx \mu^{aux} \cdot \frac{r^l}{\gamma}\left(\sqrt{1 + \frac{2\gamma}{r^l}} - 1\right) \quad [34]$$

$$GR \approx \sqrt{\frac{D^{eff}}{\alpha(r^u + r^l)}} \cdot \ln\left[\frac{r^l}{\gamma}\left(1 + \sqrt{1 + \frac{4\gamma}{r^l}}\right) + 4\right] \quad [35]$$

where $\gamma = 2\mu^{aux}/\mathcal{I}^C$. Note that for our strains we found experimentally (Fig. S7) that $\mu^{aux} = \mu^{wt}$. We can make some observations:

- The maximum growth rate does not depend on the uptake rate of amino acids but only on the leakage rate.

- The growth range (and thus the interaction range) depends strongly (square-root) on the uptake rate and the diffusion constant and weakly (logarithmic) on the leakage rate.

- When uptake rates are much higher then leakage rate ($r^l \ll r^u$), the growth range only depends on the cell density and on the ratios of uptake rate relative to diffusion ($r^u/D$) and leakage rate relative to the maximum growth rate of the auxotroph ($r^l/\mu^{aux}$).

- The cell density strongly affects the growth range by modulating the effective diffusion constant and ratio between *intra* to *extra* cellular environment (Fig. S10b).

To remind the reader: $\frac{D^{eff}}{\alpha}$ depends on the cell density $\rho$; $\rho$ affects both the diffusion constant $D^{eff} = \frac{1-\rho}{1+\rho/2} \cdot D$ and the volume ratio of *intra* to *extra* cellular environment $\alpha = \frac{\rho}{1-\rho}$. So

$$\frac{D^{eff}}{\alpha} = \frac{2(1-\rho)^2}{\rho(2+\rho)} \cdot D \qquad [36]$$

Fig. S10a shows how $D^{eff}$ and $\frac{D^{eff}}{\alpha}$ depend on density $\rho$.

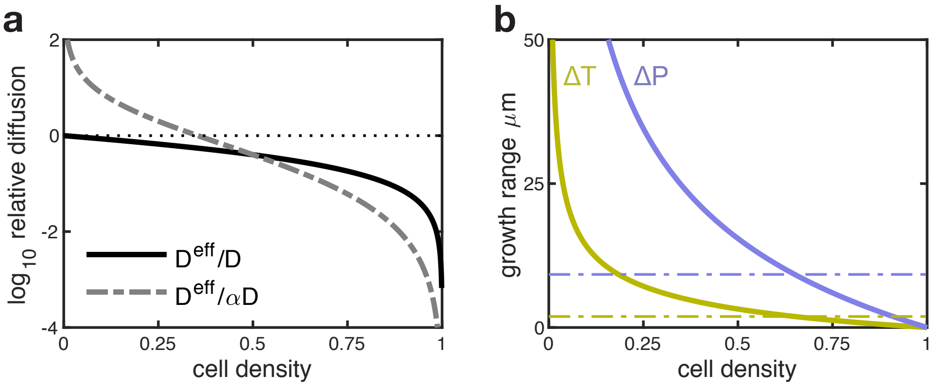

**Fig. S10. Effect of density of cells on diffusion of molecules.** (a) Dependence of $D^{eff}$ and $\frac{D^{eff}}{\alpha}$ on density $\rho$. High cellular densities reduce the effective diffusion of molecules. In our microfluidic chambers cellular density is about 0.65 (i.e. fraction of the chamber's volume occupied by cells). (b) The growth range decreases with higher densities. The analytically predicted growth range (eq. 33) is shown as function of cellular density. The dashed lines indicate the predicted growth range at the experimentally measured density. All other parameters as indicated in table S1.

### 4. Supplementary Methods

**4.1. Proportionality of analytical growth range and interaction range.** Given a set of parameters, we calculated analytically the growth range using Equation 33 and we estimated the interaction range with the model. The interaction range was estimated as follows: we ran our model on experimentally observed spatial arrangements after downscaling the segmented images to a 40x40 grid. We used the model predicted growth rates and repeated the correlation analysis described in

Methods to extract the predicted interaction range. Fig. 4b (proportionality between growth range and interaction range) is made by changing the uptake of the amino acids and keeping all other parameters fixed (see Table S1).

**4.2. Growth rate measurements in batch cultures.** Overnight cultures were started from a single colony and were grown overnight at 37°C in a shaker incubator in standard M9 glucose medium (0.2% glucose) supplemented with 50 $\mu$g/L of L-proline and 20 $\mu$g/L L-tryptophan. Subsequently, cells were inoculated in a 96-well plate in M9 glucose media without any amino acids (M9) or in M9 glucose media supplemented with 50 $\mu$g/L of L-proline (M9-P), 20 $\mu$g/L L-tryptophan (M9-T), or both (M9-PT). Cells were grown at 37°C in a plate reader and the optical density at 600nm (OD600) was measured every 3 minutes. Each strain was measured in 12 replicates. Growth rates were determined by performing a linear fit on the $\log_2$ transformed growth curves. The fit was done over a moving time window of 30min and the growth rate of each strain was determined as the maximum slope obtained over all fits.

### 5. Supplementary Tables

We list here all parameters of the individual-based model with their source.

| Parameter | Description | Value | Source |
|---|---|---|---|
| $r_1^u$ | uptake of proline | 2.04 $1/s$ | Literature[4] |
| $r_2^u$ | uptake of tryptophan | 24.05 $1/s$ | Literature[5] |
| $D_1$ | diffusion of proline | $8.79 \cdot 10^2$ $\mu m^2/s$ | Literature[6] |
| $D_2$ | diffusion of tryptophan | $6.59 \cdot 10^2$ $\mu m^2/s$ | Literature[7] |
| $\mu^{wt}$ | growth on M9 media + 0.2% glucose | 1.29 $1/h$ | Measured |
| $r_1^l$ | leakage proline | $1.59 \cdot 10^{-5}$ $1/s$ | Fitted (see section 3.4.2) |
| $r_2^l$ | leakage tryptophan | $6.04 \cdot 10^{-7}$ $1/s$ | Fitted (see section 3.4.2) |
| $\rho$ | density of cells | 0.65 | Measured |
| $dX$ | grid (cell) size | 1.5 $\mu m$ | Estimated from number of cells per chamber |

**Table S1. Parameters of individual-based model. All parameters of the model are taken from literature or measured, apart from the two leakage rates, which are estimated as described in section 3.4.2**

### 6. Supplementary Video - Caption

**Supplementary Video S1**: The 3D rendering shows the microfluidic device utilised for all experiments. The microfluidic device consists of chambers of 60x60 $\mu m$ and 0.76 $\mu m$ in height facing a feeding channel of 22 $\mu m$ in height and 100 $\mu m$ in width. *Escherichia coli* cells grow in monolayer communities in the chambers and are imaged using time-lapse microscopy. Thin tubing connects the inlet of the feeding channel to syringes containing fresh media, and the outlet to a waste collection. Media continuously flows in the feeding channel providing the bacteria with fresh nutrients as they grow in the chambers.

**Supplementary Video S2**: Two auxotrophic strains of *Escherichia coli* grow in microfluidic chambers of 60x60 $\mu m$. Left: False coloured fluorescence images show proline auxotrophic cells in purple and tryptophan auxotrophic cells in yellow. Right: The same cells are coloured based on their growth rate, with lighter colours indicating higher growth rates. Growth rates are higher for auxotrophic cells close to the partner. One cell type (in purple) has higher growth rates further away from the partner.

### 7. References

1. Sander, T., Farke, N., Diehl, C., Kuntz, M., Glatter, T., Link, H., Allosteric Feedback Inhibition Enables Robust Amino Acid Biosynthesis in E. coli by Enforcing Enzyme Overabundance. *Cell Systems.* **8**:66-75 (2019)

2. Reznik, E., Christodoulou, D., Goldford, J. E., Briars, E., Sauer, U., Segre', D., Noor, E., Genome-Scale Architecture of Small Molecule Regulatory Networks and the Fundamental Trade-Off between Regulation and Enzymatic Activity. *Cell Reports.* **20**:2666-2677 (2017)

3. Kovarova-Kovar, K. & Egli, T., Growth Kinetics of Suspended Microbial Cells: From Single-Substrate-Controlled Growth to Mixed-Substrate Kinetics. *Microbiol. Mol. Biol. Rev.* **2**:646-666 (1998)

4. Grothe, S., Krogsrud, R. L., McClellan, D. J., Milner, J. L. & Wood, J. M. Proline Transport and Osmotic Stress Response in Escherichia coli K-12. *Journal of Bacteriology.* **166**:253-259 (1986)

5. Piperno, J. R., Oxender, D. L. Amino Acid Transport Systems in Escherichia coli. *Journal. Biol. Chem.* **243**:5914-5920 (1968)

6. Wu, Y., Ma, P., Liu, Y. & Li, S. Diffusion coefficients of l-proline, l-threonine and l-arginine in aqueous solutions at 25°C. *Fluid Phase Equilibria.* 186:27-38 (2001)

7. Longsworth, L. G. Diffusion Measurements, at 25°C, of Aqueous Solutions of Amino Acids, Peptides and Sugars. *Contrib. from Lab. Rockefeller Inst. Med.* Res. Nov. 20 (1953)
